# Generation, quality control, and analysis of the first genomically humanised knock-in mice for the ALS/FTD genes *SOD1, TARDBP* (TDP-43), and *FUS*

**DOI:** 10.1101/2021.07.05.451113

**Authors:** Anny Devoy, Georgia Price, Francesca De Giorgio, Rosie Bunton-Stasyshyn, David Thompson, Samanta Gasco, Alasdair Allan, Gemma F. Codner, Remya R. Nair, Charlotte Tibbit, Ross McLeod, Zeinab Ali, Judith Noda, Alessandro Marrero-Gagliardi, José M Brito-Armas, Michelle Simon, Edward O’Neill, Jackie Harrison, Gemma Atkins, Silvia Corrochano, Michelle Stewart, Lydia Teboul, Abraham Acevedo-Arozena, Elizabeth M.C Fisher, Thomas J. Cunningham

**Affiliations:** Department of Neuromuscular Diseases, UCL Institute of Neurology, Queen Square, London, WC1N 3BG; UK MRC Harwell Institute, Harwell Campus, Oxfordshire, OX11 0RD, UK; Research Unit, Hospital Universitario de Canarias; ITB-ULL and CIBERNED, La Laguna, 38320, Spain

**Keywords:** ALS, FTD, long-range sequencing, genome engineering, allele QC, mouse, genomic humanisation, humanisation, knock-in.

## Abstract

Amyotrophic lateral sclerosis - frontotemporal dementia spectrum disorder (ALS/FTD) is a complex neurodegenerative disease; up to 10% of cases are familial, usually arising from single dominant mutations in >30 causative genes. Transgenic mouse models that overexpress human ALS/FTD causative genes have been the preferred organism for *in vivo* modelling. However, while conferring human protein biochemistry, these overexpression models are not ideal for dosage-sensitive proteins such as TDP-43 or FUS.

We have created three next-generation *genomically humanised* knock-in mouse models for ALS/FTD research, by replacing the entire mouse coding region of *Sod1*, *Tardbp* (TDP-43) and *Fus*, with their human orthologues to preserve human protein biochemistry, with exons and introns intact to enable future modelling of coding or non-coding mutations and variants and to preserve human splice variants. In generating these mice, we have established a new-standard of quality control: we demonstrate the utility of indirect capture for enrichment of a region of interest followed by Oxford Nanopore sequencing for robustly characterising large knock-in alleles. This approach confirmed that targeting occurred at the correct locus and to map homologous recombination events. Furthermore, extensive expression data from the three lines shows that homozygous humanised animals only express human protein, at endogenous levels. Characterisation of humanised FUS animals showed that they are phenotypically normal compared to wildtype littermates throughout their lifespan.

These humanised mouse strains are critically needed for preclinical assessment of interventions, such as antisense oligonucleotides (ASOs), to modulate expression levels in patients, and will serve as templates for the addition of human ALS/FTD mutations to dissect disease pathomechanisms.

## INTRODUCTION

Amyotrophic lateral sclerosis, ALS, is a relentless and devastating neurodegenerative disease that causes the progressive death of motor neurons, resulting in spreading paralysis and death typically within 5 years from diagnosis (Brown and Al-Chalabi, 2017; Hardiman et al., 2017). The lifetime risk for developing ALS is 1 in 300 in the UK, which has an average lifespan of the early 80s (Alonso et al., 2009). ALS and frontotemporal dementia (FTD) lie on a disease spectrum with overlapping genetics, pathology, and symptoms (Abramzon et al., 2020). Most ALS occurs in mid-life but a wide-span of ages has been reported, from adolescence to old age. ALS/FTD remains incurable and essentially untreatable, with two FDA approved ALS treatments that only confer, on average, a few more months of life (Brown and Al-Chalabi, 2017; Hardiman et al., 2017).The majority of ALS is sporadic (sALS), of predominantly unknown cause, but ∼10% is familial (fALS), usually with an autosomal dominant mode of inheritance, with at least 30 possible monogenic forms described in proteins with varying function.

Mutations in Superoxide Dismutase 1 (*SOD1*) were the first to be identified in ALS patients in 1993 (Rosen et al., 1993), but after >25 years of research, we still do not know how mutations in *SOD1* or those identified in other genes lead to selective neuronal death, or what shared or distinct mechanisms are at play between different genetic forms (Mejzini et al., 2019; Taylor et al., 2016). *SOD1* mutations make up ∼20% of fALS – the second-most frequent known genetic cause (Brown and Al-Chalabi, 2017). The leading cause for ALS and FTD is a hexanucleotide repeat expansion in intron 1 of *C9orf72* (DeJesus-Hernandez et al., 2011; Renton et al., 2011), whilst mutations in several RNA binding proteins including Fused in Sarcoma (FUS) and TAR DNA Binding Protein 43 (TDP-43, encoded by the *TARDBP* gene) indicate that disruption of RNA metabolism is a key pathomechanism (Zhao et al., 2018). *FUS* mutations, occurring in <5% of fALS, may cause an unusually early onset – patients as young as 11 years of age have been described, leading to a particularly aggressive disease (Picher-Martel et al., 2020). *FUS* mutations can also (albeit more rarely) cause FTD, highlighting the continuum between the two diseases (Abramzon et al., 2020). Additionally, FUS pathological aggregates (together with aggregates of related FET-family RNA binding proteins) in the absence of mutations, have been identified in ∼10% of FTD cases (Neumann et al., 2009; Urwin et al., 2010).

Whilst mutations in *TARDBP* are also relatively rare (<5% fALS), the presence of TDP-43 pathology plays a central role in ALS. In ∼97% of all ALS cases, this normally nuclear RNA binding protein aggregates into ubiquitinated and hyperphosphorylated forms in the cytoplasm of affected cells, leading to an almost complete loss of nuclear function, affecting the normal splicing of hundreds of exons (Hardiman et al., 2017; Suk and Rousseaux, 2020). TDP-43 pathology is present in a growing number of diseases collectively known as TDP-43 proteinopathies, including ALS, ∼45% of FTD cases, and a recently described subtype of dementia in old age called limbic-predominant age-related TDP-43 encephalopathy (LATE) (de Boer et al., 2020; Hardiman et al., 2017; Nelson et al., 2019). A number of key mechanisms have been proposed for how mutations in *TARDBP* or *FUS* cause neurodegeneration, from pathological protein aggregation to disrupted RNA metabolism. Interestingly, TDP-43 pathology is not present in SOD1-ALS, suggesting that at least at the pathological level they may be caused by distinct mechanisms (Mackenzie et al., 2007).

The mouse is the mammalian organism of choice for *in vivo* modelling of the complex biology of ALS/FTD and many mouse models have been invaluable for highlighting effects of mutation (De Giorgio et al., 2019). As fALS is mainly caused by dominant mutations, transgenic mice overexpressing human ALS/FTD genes have been the most widely used models, because they confer human biochemistry of the protein of interest and can model end-stage ALS within a short timeframe. However, in all transgenic strains the exogenous DNA randomly inserts into the genome and almost always concatemerises, even with large sequences from Bacterial Artificial Chromosome (BAC) donor vectors. Thus, the exogenous DNA disrupts sequences at the insertion site, which may have phenotypic outcomes unrelated to the transgenic sequence: a study of 40 commonly used strains of transgenic mice found at least 75% had altered DNA sequence around the site of insertion, including large deletions (up to a megabase) and structural changes that in half of cases disrupt at least one coding gene (Goodwin et al., 2019). Ectopic expression at non-endogenous loci may also affect spatiotemporal expression patterns and pathogenesis in unpredictable ways unrelated to human disease.

As exogenous DNA usually inserts in multiple copies, gene-dosage is altered, which usually affects protein levels derived from the transgene. Sometimes, as in the case of the widely used SOD1 G93A transgenic model of ALS, this can be an advantage and a high protein level confers a fast phenotype, enabling researchers to study disease trajectory within the ∼5-months these animals take to reach humane endpoint. However, with multiple copies, allele instability can occur within transgene arrays, and stochastic changes in copy number can dramatically alter phenotype (Acevedo-Arozena et al., 2011; Alexander et al., 2004).

Furthermore, phenotypes may be due to overexpression, not the effects of the mutation. The RNA binding proteins TDP-43 and FUS are dosage-sensitive and phenotypes arise from even mild overexpression of the wildtype human gene (Ling et al., 2019; Mitchell et al., 2013; Wils et al., 2010; Xu et al., 2010) and even low copy transgenics have to be bred to mice with the endogenous gene knocked out, in order to keep dosage as normal as possible (Lopez-Erauskin et al., 2018). While cross-breeding to endogenous gene Knock Out (KO) mice is an added complication for transgenic models, it is necessary in order to model loss of function, which has recently gained traction as being important for pathomechanism in forms of ALS (Briese et al., 2020; Brown et al., 2021; Humphrey et al., 2020; Klim et al., 2019; Ma et al., 2021; Melamed et al., 2019; Saccon et al., 2013). Finally, transgenic models that ectopically express a wildtype human ALS gene are used as controls for transgenics that express the mutant protein; however, since transgenics integrate at random in different copy numbers, comparisons between control and mutant transgenics are confounded.

Issues of insertional mutagenesis, ectopic overexpression, allele instability, the need to investigate loss of function, gene dosage-sensitivity, and lack of precise genetic control animals, may all now be addressed by the generation of physiologically relevant Knock-In (KI) strains harbouring mutations in endogenous mouse genes. Such mice express proteins/mutations of interest at endogenous levels, modelling human disease. For ALS genes, this usually leads to mild phenotypes with late disease onset, making them good models to dissect early disease mechanisms (De Giorgio et al., 2019). We and others have made KI lines for the key ALS genes *Sod1* (Joyce et al., 2015), *Tardbp* (Fratta et al., 2018), and *Fus* (Devoy et al., 2017) – all with late onset motor neuron degeneration – that have been used for dissection of ALS disease mechanisms.

However, most KI strains express the mouse protein of interest, thus not modelling human biochemistry which can be important for proteinopathies; for example, human SOD1 protein has distinct biochemical properties associated with specific residues that are not present in mouse, but are important for SOD1 misfolding, aggregation, and inferring neuronal toxicity in the context of ALS (Crown et al., 2020; DuVal et al., 2019; Nagano et al., 2015; Perri et al., 2020). Thus, our rationale for *genomic humanisation* in mice – in which the mouse endogenous gene is replaced with the human orthologous sequence, exons and introns included, and driven by the mouse promoter – is to maintain endogenous expression levels while conferring human protein biochemistry together with human splicing patterns (Nair et al., 2019; Zhu et al., 2019).

Here, we present genomically humanised lines for *SOD1*, *TARDBP* and *FUS* (*hSOD1*, *hTARDBP*, *hFUS*). Each strain was produced by homologous recombination with large homology arms (22-150 kb) in mouse embryonic stem (ES) cells, allowing us to replace the entire mouse coding sequence with its human equivalent. The generation of these mice is technically challenging and requires extensive quality control to assess allele integrity and integration at the correct endogenous locus. We present a robust genomic pipeline using indirect capture technology to enrich for high molecular weight (HMW) DNA from the targeted loci followed by long-read sequencing that allows us to validate that our humanisation strategy occurred correctly, with precise integration of the targeting vector in the correct endogenous mouse locus. We can also map the recombination events between the vector and the mouse genome. All three human genes functionally replace their mouse orthologues leading to the expression of only human protein at endogenous levels in homozygous humanised mice. Moreover, as a proof of principle, we show extensive phenotypic and molecular characterisation of the homozygous *hFUS* mice, showing that they are phenotypically normal throughout their entire lifespan, underscoring the fact that human *FUS* can functionally replace the mouse gene throughout the ageing process. These mice will be freely available to the community for driving forward novel findings in ALS/FTD and associated therapeutics.

## DESIGN

### Humanisation strategies

For each of the three genomic humanisation KI projects, the principle goal was to achieve endogenous expression of human genes in mouse. In each case we maintained the endogenous mouse promoter to drive inserted human genes, with the transition from mouse-to-human sequence beginning at the translational start codon ATG sequence; the rationale being that it was best to maintain coupling of mouse transcriptional machinery with the mouse promoter to drive each human gene at as close to physiological levels as possible. All humanisation projects entailed KI of genomic human sequence (i.e. genomic humanisation) including all coding exons and intervening introns. Introns were included (rather than a simpler KI of cDNA) to prevent undesired disruption to physiological expression, for example to intronic autoregulatory elements (Humphrey et al., 2020). Including *human* introns serves two key purposes: first it enables modelling of known human intronic variants and mutations, and second it maintains human splicing complexity and affords study of human gene regulation. For the 3’ end of each locus, we employed a bespoke approach.

For *hSOD1*, a complex conditional allele was engineered, incorporating a duplication of exon 4, intron 4, exon 5, the human 3’ UTR, and ∼1 kb of downstream mouse terminator sequence (**Figure 1A, 1B**). In this configuration, the human gene is transcribed and translated as normal, not incorporating the downstream duplicated sequence. The upstream copy of this sequence is flanked by *loxP* sites, such that following CRE recombination, the downstream copy of the sequence (exons 4’ to 5’) is brought into frame and is expressed. This allows for conditional expression of mutations, or reversion from mutation to wildtype, of mutations placed in the downstream or upstream duplicated exons, respectively.

**Figure 1.**
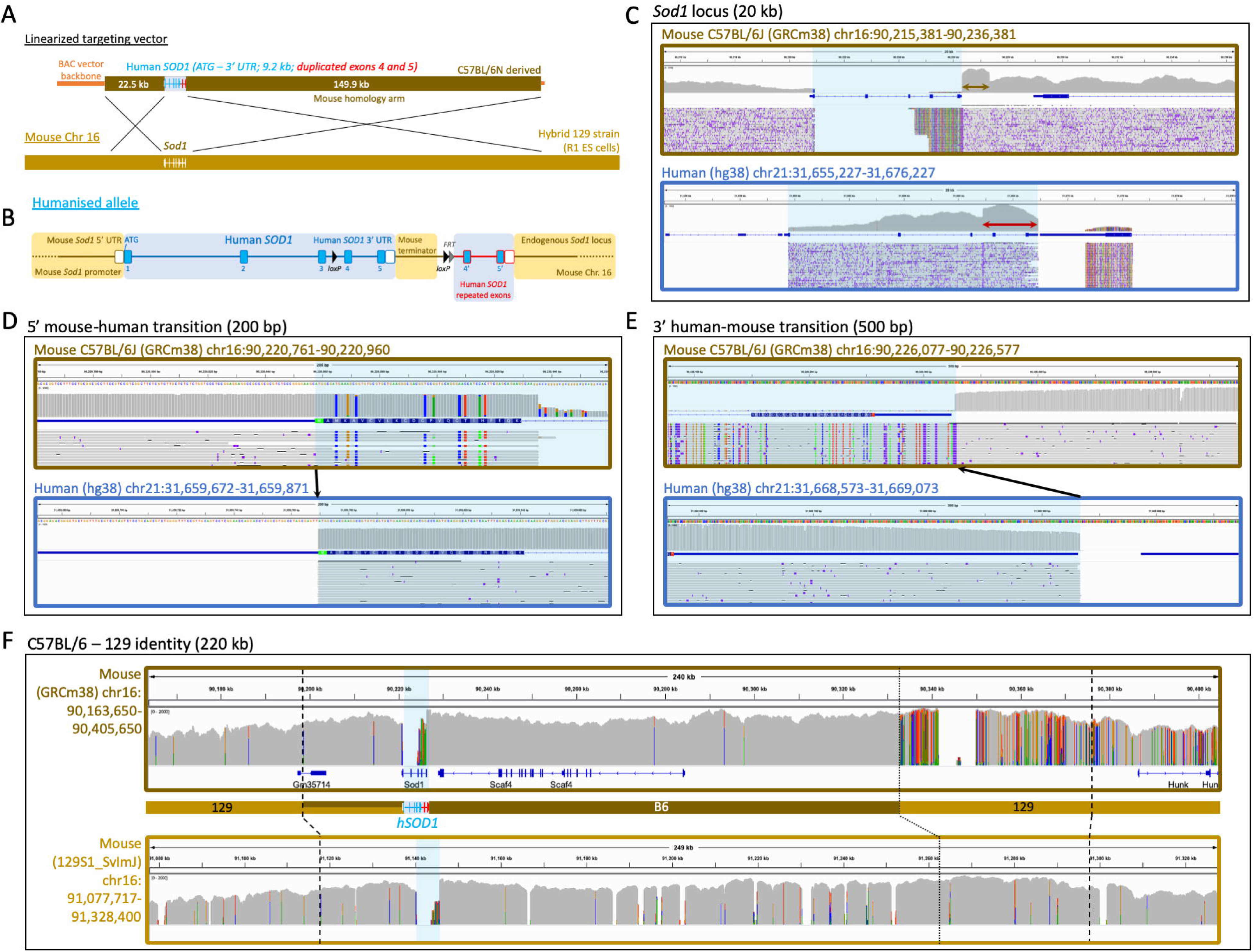
Genomic humanisation of the endogenous mouse *Sod1* gene. **(A)** A BAC construct, engineered to harbor the human *SOD1* genomic sequence (blue) flanked by large regions of mouse homology (brown), was used as a donor to replace endogenous mouse *Sod1* with human *SOD1* in mouse ES cells via homologous recombination. **(B)** The humanised *SOD1* allele in more detail, showing humanisation of *Sod1* from the ATG start codon to the end of the 3’ UTR (blue); human exon 4, intron 4, exon 5, the 3 ‘UTR, plus 1 kb of downstream mouse terminator sequence was floxed and duplicated to generate a conditional cassette. An *FRT* site downstream of the second *loxP* site is a remnant of selection cassette removal. **(C-F)** IGV visualizations of Oxford Nanopore read alignments via minimap2. Alignment of nanopore sequencing reads across **(C)** the mouse and human *SOD1* gene loci; **(D)** the mouse and human *SOD1* ATG start codon; **(E)** the mouse and human *SOD1* 3’ UTR; and **(F)** the wider *SOD1* locus comparing C57BL/6 and 129 mouse strain identity, with dashed lines representing the boundaries of the homology arm region and dotted lines delineating C57BL/6 and 129 strain genome identity. Blue shading in alignment screenshots represents the humanised region. Brown and red arrows in (C) denote the engineered duplications. Black arrows in (D) and (E) show the precise transition from mouse-to-human and human-to-mouse sequence.

For *hTARDBP*, a simpler approach was taken and humanisation extends only from the start ATG to the stop codon sequence (**Figure 2A, 2B**). As 3’ UTR disruption has been shown to affect expression of the downstream gene *Masp2* (Dib et al., 2014), which lies tail-to-tail with *Tardbp* so that both overlap in their 3’ UTRs, we chose to keep the endogenous mouse 3’ UTR sequence to maintain correct *Masp2* expression.

**Figure 2.**
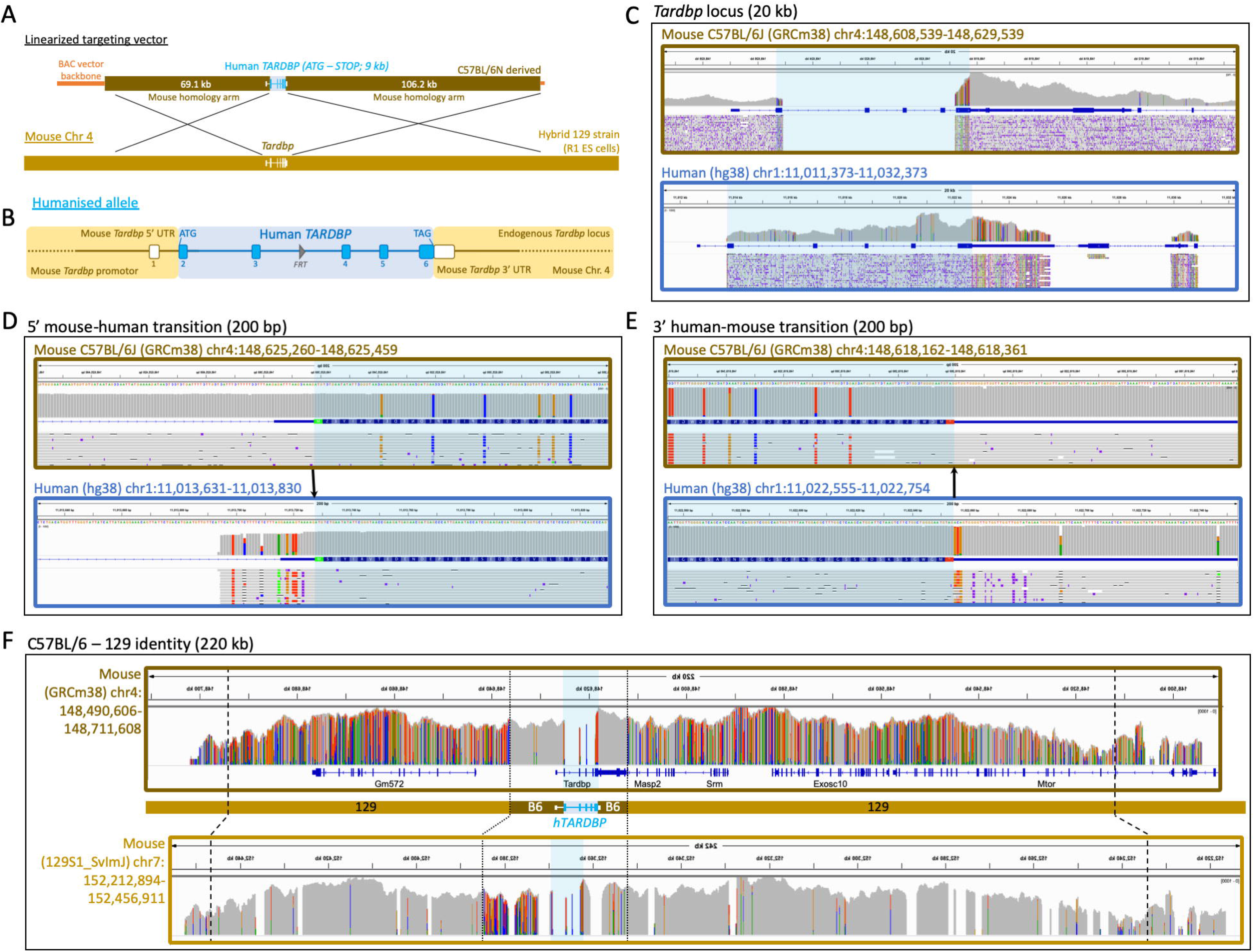
Genomic humanisation of the endogenous mouse *Tardb*p gene. **(A)** A BAC construct, engineered to harbor the human *TARDBP* genomic sequence (blue) flanked by large regions of mouse homology (brown), was used as a donor to replace endogenous mouse *Tardbp* with human *TARDBP* in mouse ES cells via homologous recombination. **(B)** The humanised *TARDBP* allele in more detail, showing humanisation of *Tardbp* from the ATG start codon to the TAG stop codon (blue). An *FRT* site in intron 3 is a remnant of selection cassette removal. **(C-F)** IGV visualizations of Oxford Nanopore read alignments via minimap2. Alignment of nanopore sequencing reads across **(C)** the mouse and human *TARDBP* gene loci; *(D)* the mouse and human *TARDBP* ATG start codon; **(E)** the mouse and human *TARDBP* stop codon; and **(F)** the wider *TARDBP* locus comparing C57BL/6 and 129 mouse strain identity, with dashed lines representing the boundaries of the homology arm region and dotted lines delineating C57BL/6 and 129 strain genome identity. Blue shading in alignment screenshots represents the humanised region. Black arrows in (D) and (E) show the precise transition from mouse-to-human and human-to-mouse sequence.

In the case of *hFUS,* the transition from human to mouse was placed after the 3’UTR (**Figure 3A, 3B**). The human 3’ UTR was included because (1) variants in the 3’ UTR have been linked to risk for developing ALS (Dini Modigliani et al., 2014; Sabatelli et al., 2013) and (2) ALS-frameshift mutations at the 3’ end of FUS incorporate sequence from the human 3’ UTR and if not humanised, such mutations would result in longer 3’ neopeptides that may impact pathogenicity (An et al., 2020). In addition, *loxP* sites were placed upstream of exon 15 and downstream of the 3’ UTR to allow for future conditional studies.

**Figure 3.**
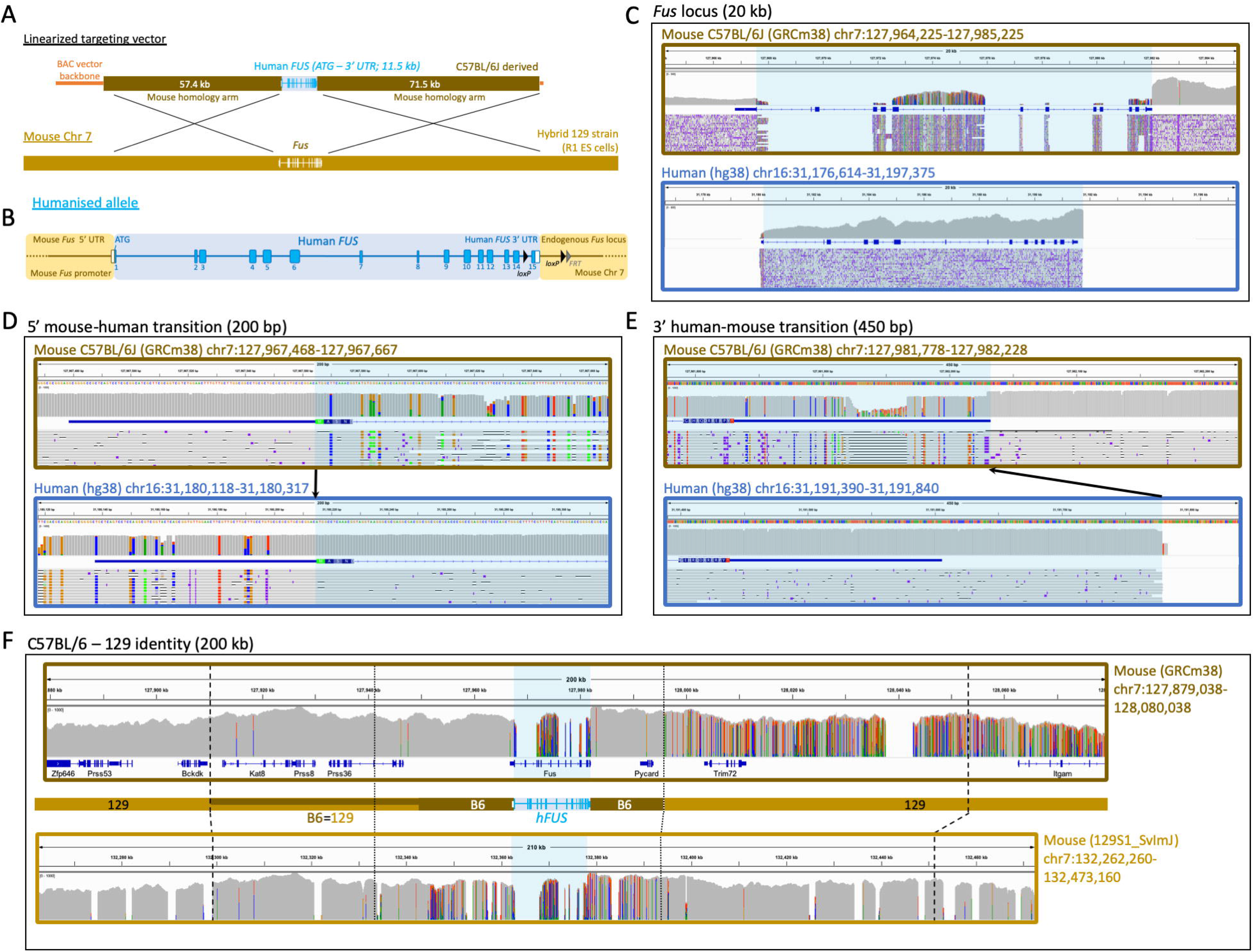
Genomic humanisation of the endogenous mouse *Fus* gene. **(A)** A BAC construct, engineered to harbor the human *FUS* genomic sequence (blue) flanked by large regions of mouse homology (brown), was used as a donor to replace endogenous mouse *Fus* with human *FUS* in mouse ES cells via homologous recombination. **(B)** The humanised *FUS* allele in more detail, showing humanisation of Fus from the ATG start codon through to the end of the 3’ UTR, including all coding exons and introns, driven by the endogenous *Fus* promoter. A *loxP/FRT* site 1 kb downstream of the 3’ UTR is a remnant of selection cassette removal. A second *loxP* site is also present in the last intron. **(C-F)** IGV visualizations of Oxford Nanopore read alignments via minimap2. Alignment of sequencing reads across **(C)** the mouse and human *FUS* gene loci; **(D)** the mouse and human *FUS* ATG start codon; **(E)** the mouse and human *FUS* 3’ UTR; and **(F)** the wider *FUS* locus comparing C57BL/6 and 129 mouse strain identity, with dashed lines representing the boundaries of the homology arm region and dotted lines delineating C57BL/6 and 129 strain genome identity. Blue shading in alignment screenshots represents the humanised region. Black arrows in (D) and (E) show the precise transition from mouse-to-human and human-to-mouse sequence.

### Validation approach for engineered alleles

Humanised targeting constructs for all three alleles were engineered from large BAC targeting vectors using recombineering technology (Copeland et al., 2001), with the removed mouse sequence ranging from 5.5 kb-14.5 kb and the KI replacement sequence (after selection cassette removal) ranging from 9-11.5 kb (**Figures 1A, 2A, 3A**). Constructs were electroporated into R1 ES cells, and correctly targeted clones (validated by quantitative PCR copy-count assays, data not shown) were karyotyped and injected into donor blastocysts for mouse production. Considering the relatively large size of each KI, and homology arms used in gene targeting (together spanning between 140-184 kb for each allele), standard techniques such as long-range PCR, Sanger sequencing, or Southern blotting were insufficient to validate allele integrity. To be certain of structural and sequence-level allele integrity, at the correct locus, we reasoned that high coverage long-read sequencing data was required. Therefore, we sought to establish the utility of Xdrop (Samplix) indirect target locus enrichment (Blondal et al., 2021) followed by Oxford Nanopore sequencing for large, complex allele validation. As further validation, and to corroborate DNA sequencing data, we analysed mRNA and protein expression in several tissues in all three lines. In addition, for the *hFUS* line, we took validation a step further and performed both transcriptomic and behavioural analyses.

## RESULTS

### Xdrop indirect target locus enrichment and Nanopore sequencing to confirm allele integrity in humanised mice

For long-read sequencing allele validation we first sought to establish mice homozygous for each allele. All three new strains are viable and healthy in homozygosity, with progeny produced with the expected Mendelian ratios (data not shown). Cohorts for all analyses, including homozygotes for sequencing, were produced by intercrossing heterozygous humanised mice, maintained on a C57BL/6J background, with the selection cassette removed for h*SOD1* and h*FUS*, but still present for the more recently established *TARDBP* line.

High molecular weight genomic DNA from homozygous humanised animals was prepared for indirect target locus enrichment using Xdrop™ technology (Samplix) followed by Oxford Nanopore sequencing (Blondal et al., 2021). Briefly, this approach encapsulates high molecular weight DNA into droplets, followed by droplet PCR using detection primers for the loci of interest facilitating FACS sorting of droplets containing the desired loci. Subsequently, droplet-based multiple displacement amplification provides sufficient DNA for Nanopore sequencing with high coverage (**Figure S1**). For each humanisation project, we designed between 3 and 4 pairs of detection primer sequences (**Table S1**) across each respective mouse target loci, used to enrich genomic DNA from the surrounding locus, reaching beyond the extent of the long homology arms (>20 kb) to scrutinise any potential undesired recombination events. This was successfully achieved, with an island of coverage spanning between 200-300kb for each locus, with coverage peaking around detection sequences (ranging from 50x to >1000x) (**Figures S2,S3,S4**).

Alignment (Minimap2 (Li, 2018)) of reads to the C57BL/6J (GRCm38.p6) reference genome across each targeted gene showed, in each case, a loss of the targeted mouse gene; while alignment to the human (hg38) reference genome showed an island of coverage representing the inserted human sequence (**Figures 1C, 2C, 3C**). While two of the lines (*hSOD1* and *hTARDBP*) utilised C57BL/6N-derived targeting constructs, we did not align to the C57BL/6N reference due to lack of differences between C57BL/6J and C57BL/6N at these loci, but also because the C57BL/6J reference is the most complete available, with no gaps. The immediate sequence surrounding the mouse-human and human-mouse boundaries at the 5’ and 3’ ends of the loci has sufficient sequence conservation to align to the opposing species alignment, which revealed the expected divergences (coloured lines) from each reference genome, and showed that the intended, precise transitions have been incorporated into the genome (**Figures 1D,1E,2D,2E,3D,3E**). The intended precise insertion can also be demonstrated by alignment of reads to a sequence file of the expected engineered allele (**Figures S5,S6,S7**). This revealed uninterrupted alignment profiles, in contrast to mouse and human reference alignments, with engineered insertions including *loxP* and *FRT* sites clearly visible in the alignment tracks (**S5,S6,S7**).

Large engineered structural variants in the *hSOD1* allele – duplication of the 3’ human sequence, plus duplication of the mouse terminator sequence - were clearly visible as sharp 2-fold steps in coverage in the profiles of human and mouse alignments; whilst alignment to the expected allele resolved these coverage anomalies (***Figure 1C; Figure S5***).

Two anomalies in the alignment profile against the human reference genome, in introns 2 and 5 of the *hTARDBP* allele, respectively, warranted further investigation (**Figure S6; red arrows, region X and Y**). Each region exhibited a concentration of base positions with high error rates, although at each position, the correct base was called in the majority (>70% for each affected nucleotide). Additionally, region Y in intron 5 exhibited an anomalous step-up in coverage. Both regions map precisely to the boundaries of highly repetitive SINE elements (**Figure S8**); we suspected mapping error as the majority of reads mapped correctly, many of which mapped beyond the boundaries of these features on both sides. Alignment with alternative software, NGMLR (Sedlazeck et al., 2018), built specifically to correctly identify structural variants (versus minimap2, built for fast alignment), resolved the alignment anomalies (**Figure S6**). We also PCR amplified and Sanger sequenced intron 5 using various primer sets, observed the expected band sizes, and found no evidence of sequence anomalies corresponding to the nucleotide positions with high error rates via Nanopore sequencing, or in any other position (data not shown).

Alignment of reads to the C57BL/6J reference genome across the wider targeted locus for each mouse line showed that while the human insertion and proximal-most regions align as expected, more distal homology arm correspondent regions have evidence of misalignment (**Figure 1F,2F,3F; top panels**). Our strategy used 129 strain ES cells with targeting vector homology arms derived from the C57BL/6J or C57BL/6N (herein referred to C57BL/6) genetic background, followed by backcrossing all to C57BL/6J, and so we additionally aligned reads from each mouse to the available 129S1_SvImJ reference genome for comparison **(Figure 1F,2F,3F; bottom panels**). This demonstrated the transitions from C57BL/6 identity to 129 strain identity (where SNPs polymorphic for both strains exist) and thus pinpointed the homologous recombination breakpoints (i.e. the extent of flanking sequence from the targeting vector that recombined into the respective loci together with the humanised genes). This is most clearly demonstrated in the *hTARDBP* allele, where C57BL/6 and 129 strains are divergent on both homology arms (**Figure 3F**). 11 kb (5’ to *hTARDBP*) and 5.5 kb (3’ to *hTARDBP*) of the proximal homology arm correspondent regions are derived from the C57BL/6 targeting vector, whilst the more distal homology arm correspondent regions are clearly 129 derived, showing that only a minor fraction (16.5/175 kb total) of the homology arms recombined into the locus together with the *hTARDBP* gene (**Figures S9,S10**).

For the *hFUS* allele, ∼30 kb of the proximal 5’ homology arm correspondent region is clearly derived from the C57BL/6 targeting vector, while the remaining distal 27 kb of the 5’ homology arm has no appreciable divergence between strains to make a clear determination (**Figure S11**). For the *hFUS* 3’ homology arm region, the proximal 15 kb is C57BL/6 derived with the exception of a single SNP that maps to the 129 strain, suggesting that recombination did not occur cleanly at a single point (**Figure S12**). The distal 56.5 kb of the *hFUS* 3’ homology arm region is clearly 129 strain derived. For *hSOD1*, the relatively shorter 22.5 kb 5’ homology arm region is conserved between C57BL/6 and 129 strains, whilst in the 3’ homology arm region where divergent sequence exists, albeit sparsely, the homologous recombination breakpoint is ∼100 kb from *hSOD1* with the distal 50 kb clearly mapping to 129 strain (**Figure S13**).

In all three humanisation projects we observe seamless alignment stretching beyond the limit of the homology arm regions and, together with the clear 129-C57BL/6 transitions that show recombination breakpoints, we can be confident the integrations occurred at the correct loci.

After accounting for the humanisation event, engineered insertions, the anomalies in *hTARDBP* introns 2 and 5, and strain differences, very few minor alignment imperfections remain. A number of gaps (of varying sizes) lie within the 129 strain alignments, including where the sequence is evidently of 129 strain identity. These all map to gaps in the 129 Reference Genome (sequence denoted by strings of Ns) rather than any untoward misalignment. In addition, there a number of known (human population) SNPs present in the human gene sequence in the *hSOD1* and *hTARDBP* targeting vectors, all within non-coding regions, which differ from the respective reference human and mouse genomes as reflected in the alignments (marked by **#; Figures S5,S6**). Finally, we saw positions in which individual nucleotides were flagged (coloured lines) because they had an increased error rate in the sequencing with respect to the reference sequence. Such positions are all in non-coding regions, are highly polymorphic, and map to homopolymer sequences, low complexity sequences, and repetitive element regions, which are a known cause for higher error rates in long-read sequencing data.

### Humanised mouse strains express human, not mouse, gene products

To complement the long-read sequencing data, we next sought to confirm the expression of human and not mouse gene products in the three lines. Starting with RT-PCR, we designed primers specific to each of the mouse cDNA sequences and primers specific to the human cDNA sequences to specifically test for the loss of mouse and gain of human mRNA in each respective line. In all three lines, we observed loss of mouse mRNA in homozygous humanised animals, absence of human mRNA in wildtype littermates, and gain of human mRNA in heterozygous and homozygous humanised animals (**Figures 4A,5A,6A**).

**Figure 4.**
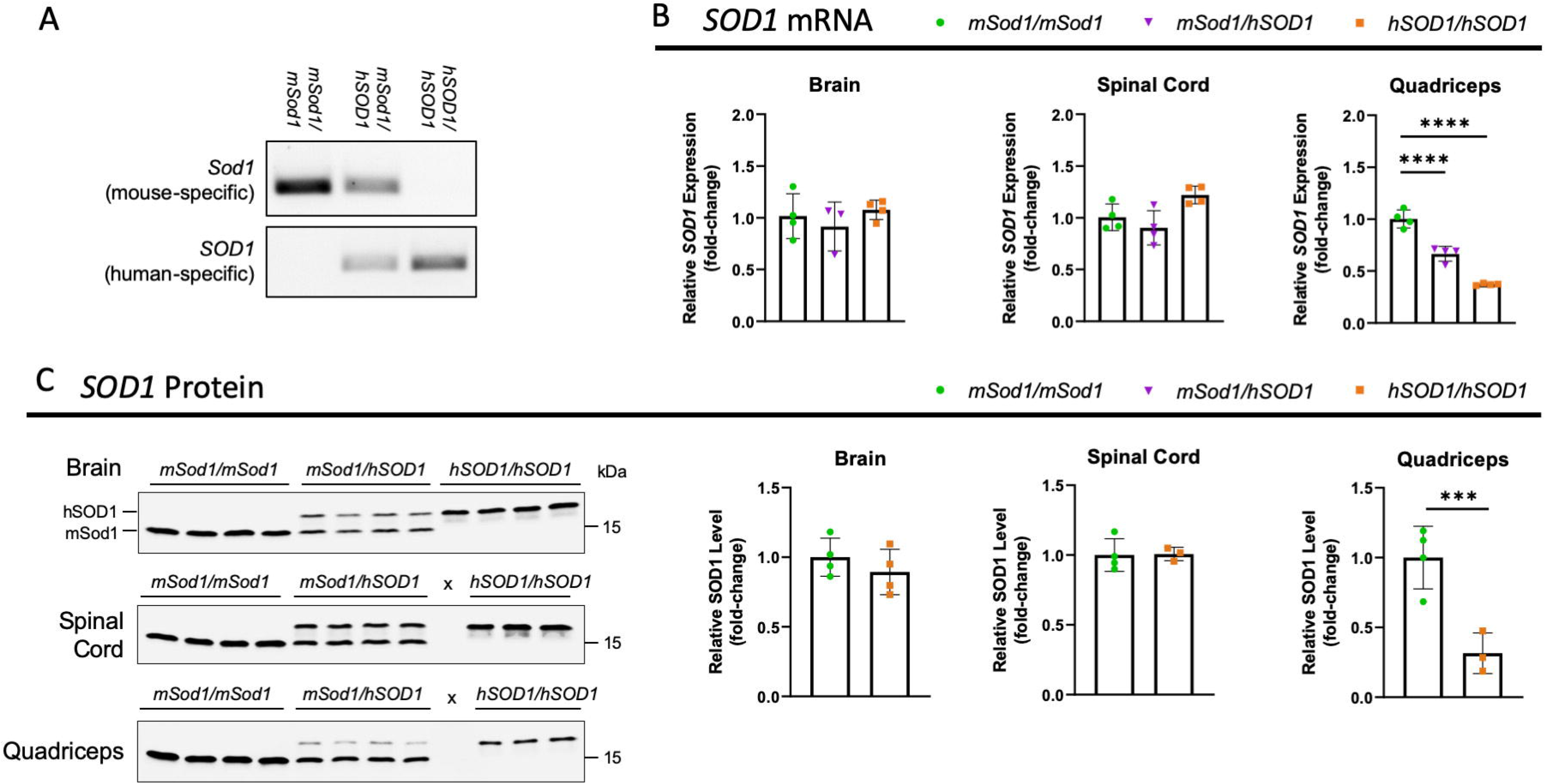
*hSOD1/hSOD1* mice only express human *SOD1*. **(A)** RT-PCR in brain from 4-month old male mice using mouse-specific *Sod1* and human-specific *SOD1* primers. **(B)** Quantitative PCR using conserved mouse-human *Sod1*/*SOD1* primers in tissues from male 4-month old mice (n=3-4 per genotype). Mean ± SD, One-way ANOVA with Dunnett’s post hoc test. **(C)** Immunoblots using a pan-mouse-human SOD1 antibody in tissues from male 4-month old mice (n=3-4 per genotype). Human SOD1 has lower mobility than mouse Sod1 in SDS-PAGE and appears as a higher band. Heterozygote quantification not included for protein data due to confounding issue of double bands. Blots normalized to total protein (**Figure S16**). Unused lanes marked by x. Mean ± SD, Unpaired t-test. \**P* < 0.05, \*\*\**P* < 0.001, \*\*\*\**P* < 0.0001.

**Figure 5.**
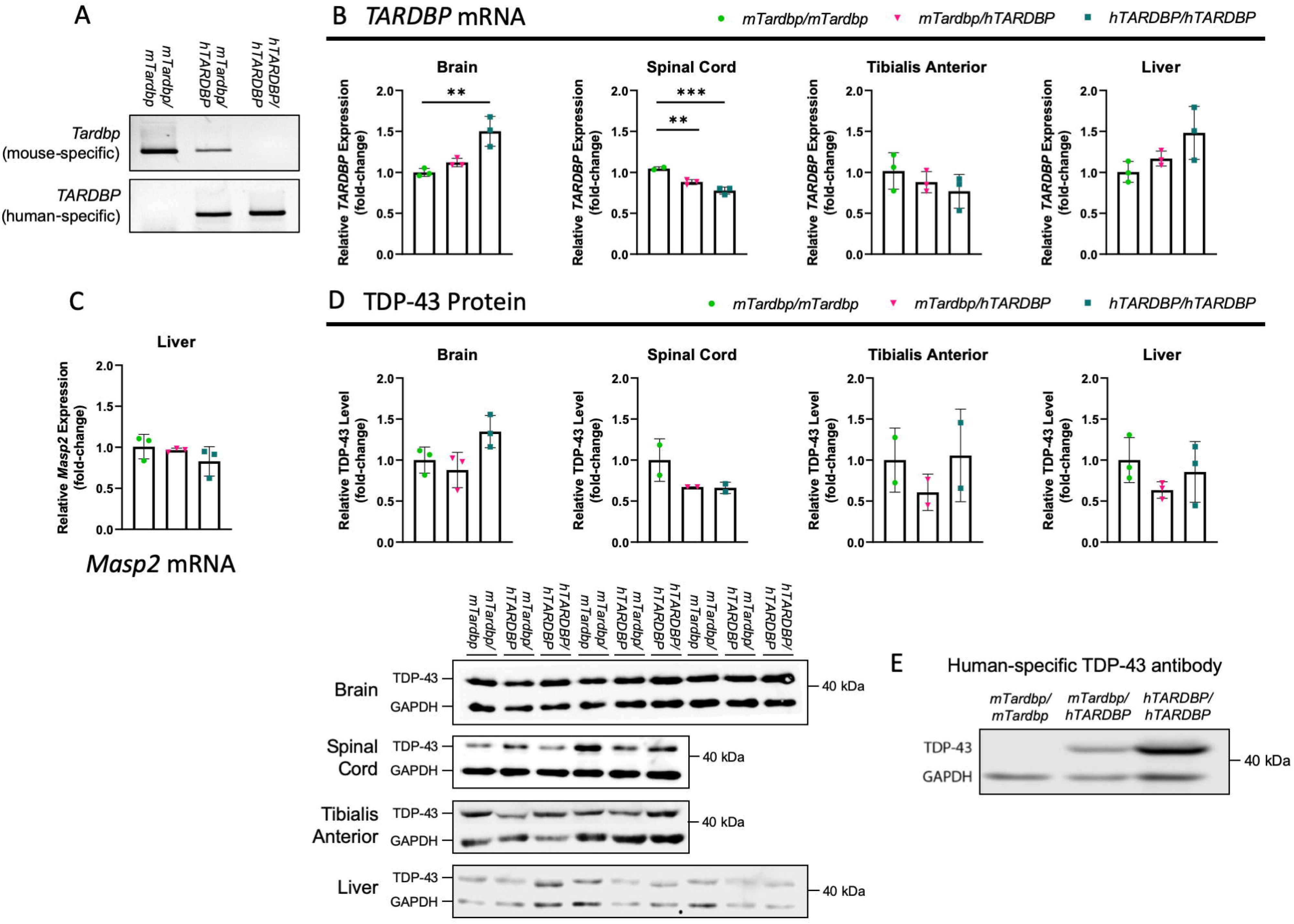
*hTARDBP/hTARDBP* mice only express human *TARDBP*. All tissues analysed are from female 14-week old *hTARDBP* mice and controls. **(A)** RT-PCR in brain using mouse-specific *Tardbp* and human-specific *TARDBP* primers. **(B)** Quantitative PCR using conserved mouse-human *Tardbp*/*TARDBP* primers (n=2-3 per genotype per tissue). **(C)** Quantitative PCR using mouse *Masp2* primers in liver (n=3 per genotype). **(D)** Immunoblots using pan-mouse-human TDP-43 antibody (n=2-3 per genotype). **(E)** Immunoblot using human-specific TDP-43 antibody in brain. \*\**P* < 0.01, \*\*\**P* < 0.001, Mean ± SD, One-way ANOVA with Dunnett’s post hoc test.

**Figure 6.**
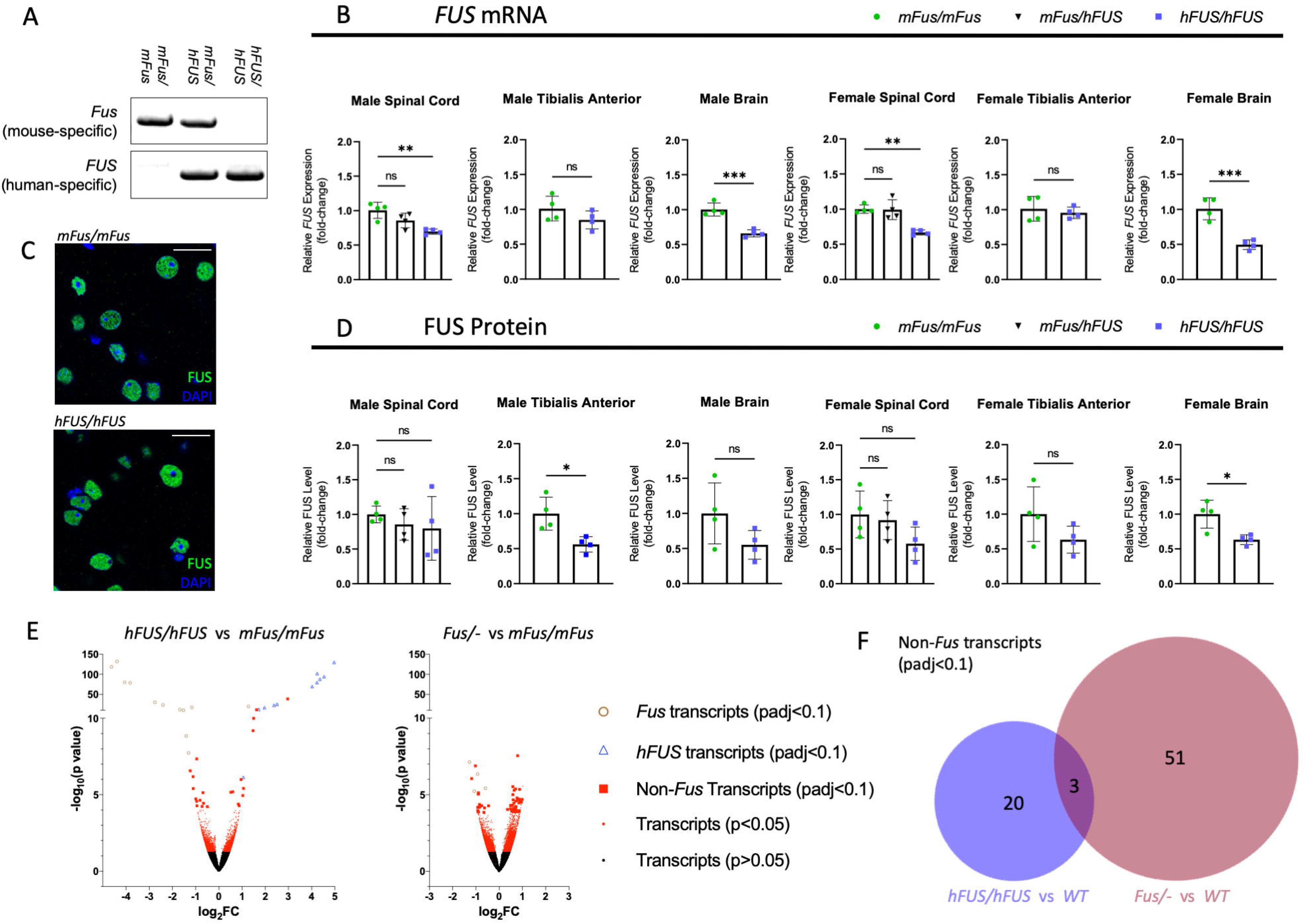
*hFUS/hFUS* mice only express human *FUS* but have minimal transcriptomic disruption. **(A)** RT-PCR from E14.5 embryonic brain using mouse-specific *Fus*, and human-specific *FUS* primers. **(B)** Quantitative RT-PCR using conserved mouse-human *Fus*/*FUS* primers in tissues from male and female 18-month old mice (n=4 per genotype per sex). Data presented as mean ± SD, ns=not statistically significant, **=p≤0.01, ***=p ≤0.001 calculated using One-way ANOVA with Dunnett’s post hoc test or Student’s T test. **(C)** Immunofluorescent staining of FUS (green) in 8-month old motor cortex showing nuclear localisation of FUS. Scale bar = 20 µm. **(D)** Quantification of immunoblots using pan-mouse-human FUS antibody in male and female spinal cord, brain, and tibialis anterior (TA) muscle tissue collected from mice aged 18 months (n=4 per genotype per sex). Blots normalized to GAPDH (**Figure S16**). Data presented as mean ± SD, ns=not statistically significant, *=p≤0.05 calculated using One-way ANOVA with Dunnett’s post hoc test or Student’s T test. Heterozygotes only included for spinal cord data. **(E)** RNAseq volcano plots showing the statistical distribution of differentially expressed transcripts between *hFUS*/*hFUS* and *mFus/mFus* (left) and *mFus/-* and *mFus/mFus* (right), from 14-week old spinal cord tissue (n=4 per genotype). Significant (padj<0.1) mouse *Fus* transcripts are shown by brown open circles, significant human *FUS* transcripts are shown by blue open triangles, significant non-*Fus* transcripts are shown by red squares, and all other transcripts are shown by red dots (p<0.05) and black dots (p>0.05). *(F)* Venn diagram comparing the number of significant non-*FUS* differentially expressed transcripts between *hFUS/hFUS* and *WT* (blue; left) and *mFus*/- and *mFus/mFus* (red; right).

Quantitative RT-PCR (qRT-PCR) and complementary immunoblots were then performed from multiple tissue types to quantify expression of the human gene products. Starting with *hSOD1* animals, mRNA and protein levels in the central nervous system (CNS; brain and spinal cord), were not significantly different at 4 months (**Figure 4B,C**), suggesting that the insertion of the conditional allele does not affect expression *per se*. In contrast, expression in muscle was significantly reduced at both the mRNA and protein level compared to controls (**Figure 4B,C**), dipping below 50% of wildtype levels, suggesting an interesting muscle-specific alteration in gene regulation that warrants future investigation. Mouse SOD1 protein has evident reduced mobility versus human SOD1, so we were able to further corroborate loss of mouse and gain of human SOD1 expression in homozygous humanised mice (**Figure 4C**).

In *hTARDBP* animals, at 14 weeks of age, we see minor differences in *Tardbp/TARDBP* mRNA expression levels between homozygous humanised mice and controls in brain and spinal cord, but not in skeletal muscle or liver (**Figure 5B**). These are not reflected in any significant changes at the TDP-43 protein level in any of the four tissues analysed (**Figure 5D**). To further assess TDP-43 functionality, we assessed the splicing of two cassette exons known to be controlled by TDP-43 function (exon 18 of *Sortilin* and exon 5 of *Eif4h*, (Fratta et al., 2018)) by RT-PCR from brain and spinal cord and found no differences in exon inclusion between any of the genotypes (**Figure S14**), suggesting that human TDP-43 protein can functionally replace the mouse protein splicing function. To verify that human TDP-43 protein is indeed expressed in *hTARDBP* mice, we took advantage of a monoclonal antibody that recognises human but not mouse TDP-43. As expected, the human-specific antibody only recognised TDP-43 in homozygous and, to a lesser extent, in heterozygous humanised mice (**Figure 5E**). As mouse *Tardbp* and *Masp2* genes are located tail-to-tail and share their 3’UTR, we assessed *Masp2* expression in liver, where it is mainly expressed, and found that, as expected, its expression was not affected by the humanisation strategy leaving the mouse 3’UTR intact (**Figure 5C**).

For *hFUS* animals, we were able characterise expression of the humanised gene more extensively in males and females during ageing. At 18 months, we observed non-significant to mild reductions in mRNA and protein expression in *hFUS* spinal cord, brain, and TA muscle versus wildtype (**Figure 6B,D**); a finding mirrored at an earlier 14 week timepoint (**Figure S15**). Mouse FUS protein has fewer amino acid residues versus human FUS (518 versus 526); this is reflected in the immunoblots where band sizes in humanised sample tissue run marginally more slowly than wildtype (**Figure S16**). In addition, we were able to show that hFUS protein correctly localises to the nucleus (**Figure 6C**); in contrast to localisation in FUS-ALS and FUS-FTD patients, where FUS protein is found to mislocalise to the cytoplasm (Kwiatkowski et al., 2009; Urwin et al., 2010; Vance et al., 2009).

Taken together, while we do observe alterations in expression that may be driven by changes in transcriptional regulation, levels are maintained within a physiological range, especially within the central nervous system.

### Homozygous *hFUS* mice display minimal transcriptomic disruption

Since FUS is an RNA binding protein and regulates RNA metabolism on a genome wide scale, we sought to gauge the transcriptomic impact of humanising mouse *Fus*. We conducted RNAseq of spinal cord tissue at 14 weeks of age, comparing *hFUS* homozygous mice with wildtype littermates. We included heterozygous KO (*mFus*/-) animals as a measure of mild loss of function. We included human *FUS* transcripts together with the mouse reference assembly when mapping reads; as expected, we saw expression of *hFUS* transcripts and concomitant loss of mouse *Fus* transcripts in *hFUS* animals (**Figure 6E**, blue triangles and brown circles; **Table 1**). Taking aside *FUS*/*Fus*-related transcripts, very few transcripts were differentially expressed in *hFUS* animals; and fewer in comparison to *mFus/-* spinal cord (**Figure 6E,F; Tables 1 and 2**). Only three non-*Fus* transcripts (Hk1-220, Arid1b-201, and Fyttd1-213) were differentially expressed in both *hFUS* homozygous and *mFus/-* spinal cord, versus wildtype; with expression of all three changing in the same direction, suggesting a mild loss of function effect, at least for these three transcripts, in *hFUS* homozygotes. Amongst the 23 non-*Fus* related differentially expressed transcripts (including 2 *Fus* pseudogenes with strong downregulation that we presume to be misaligned with *Fus*) found to be statistically significantly differentially expressed in *hFUS* homozygous spinal cord tissue, 13 are on chromosome 7. These 13 transcripts are sparsely spread across a 30.9 Mb chromosomal region surrounding the *Fus* locus, but not including genes in the immediate vicinity of the *Fus* locus (the closest affected genes are *Gdpd3* and *Tial1*, 1.2 Mb and 472 kb away, respectively). Considering that 129 strain ES cells were used to generate these mice, followed by backcrossing to the C57BL/6J genetic background, we expected a large chromosomal region surrounding the *Fus* locus to remain 129 strain identity through genetic linkage to the targeted locus for which we are selectively breeding. Indeed, whole genome SNP analysis of *hFUS* mice (of the same generation as RNAseq animals) revealed that 46 SNPs of 129 identity (specifically the 129X1/SvJ strain) clustered within a 12.3 Mb span (chr7: 117044155-129357839) of the locus surrounding *hFUS*, encompassing 10 of the 13 differentially expressed non-*Fus* transcripts on chromosome 7 (**Data file S1**). This SNP data strongly suggests differentially expressed transcripts on chromosome 7 are primarily due to sequence differences between 129 and C57BL/6J strains, reducing the direct overall transcriptomic impact of the humanisation of *Fus;* while also providing even further corroboration that the targeted locus lies in the expected region on chromosome 7.

**Table 1.**
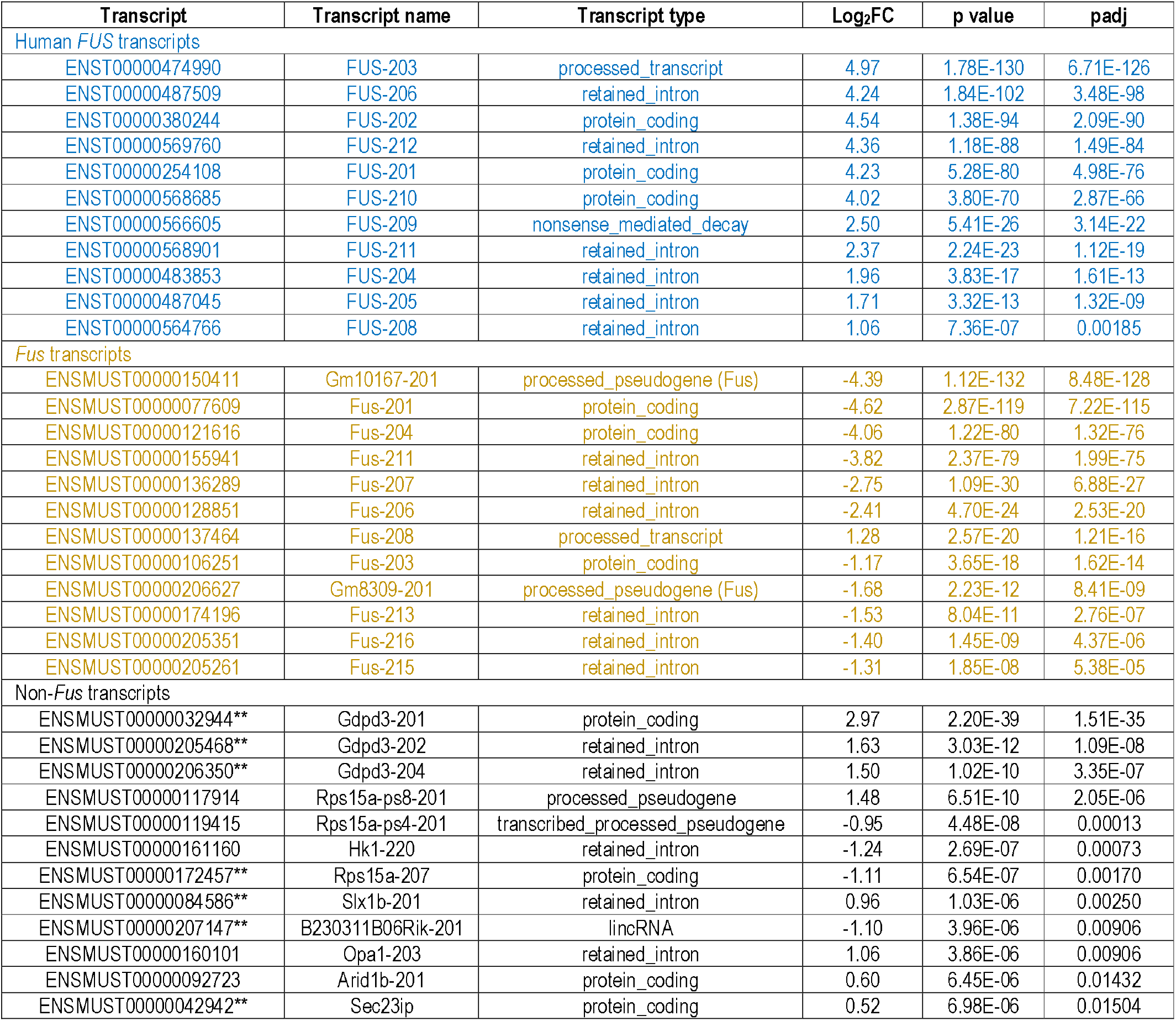
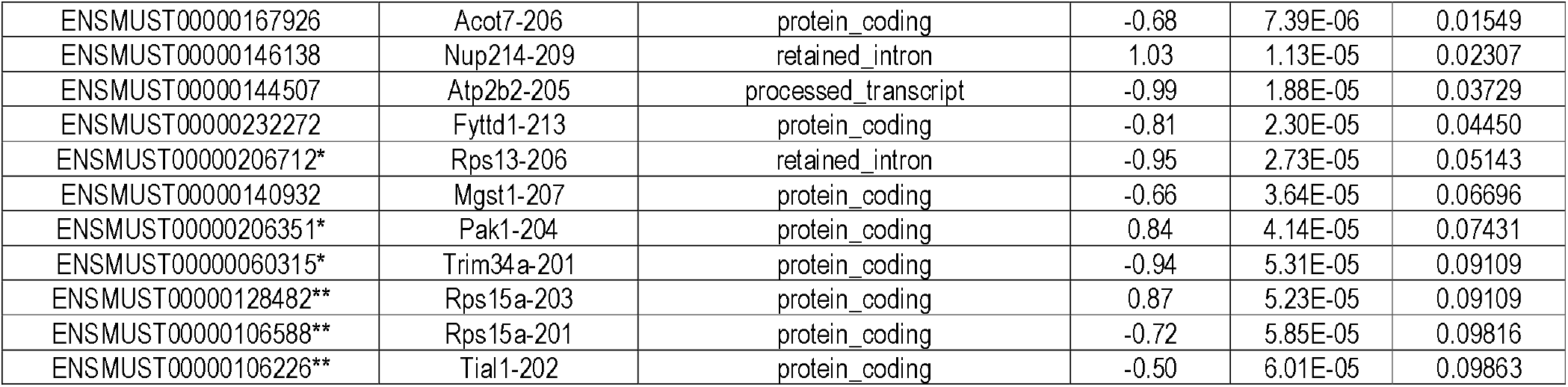
List of differentially expressed transcripts in *hFUS/hFUS versus mFus/mFUS* mice.* indicates non-Fus transcripts on chromosome 7, ** indicates non-*<u>Fus</u>* transcripts in a region of preserved 129 identity surrounding the *Fus* locus.

**Table 2.**
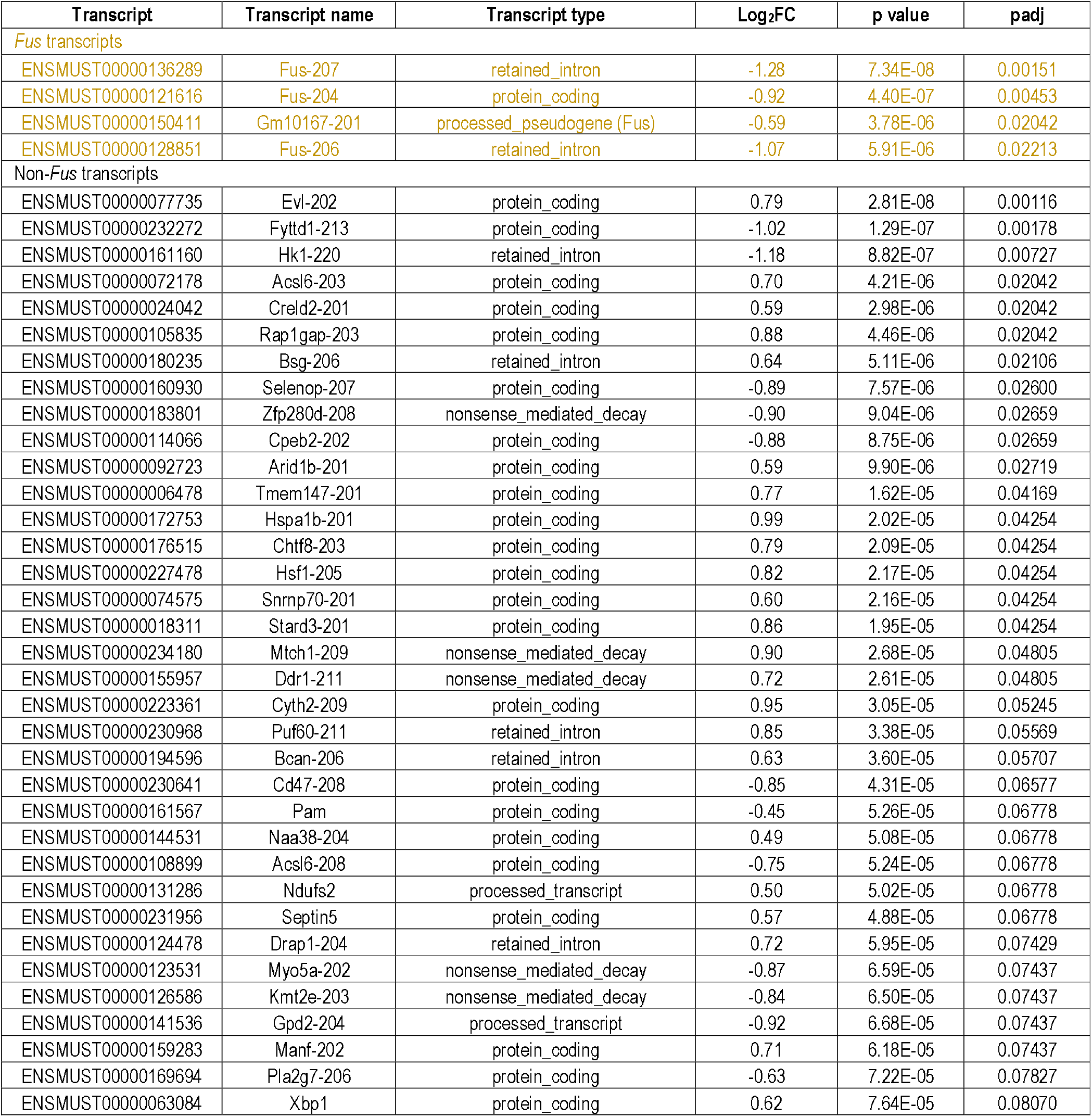
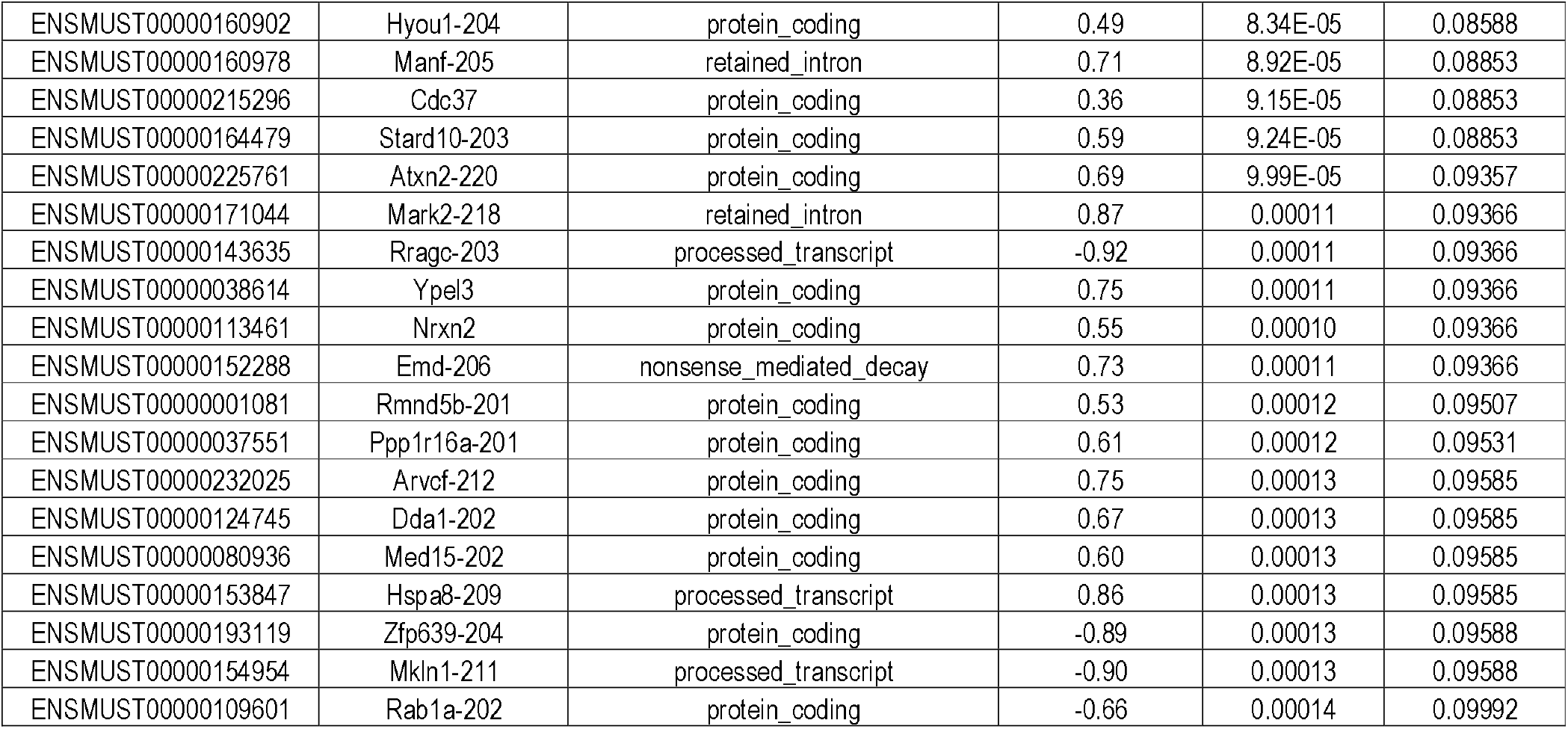
List of differentially expressed transcripts in *mFus*/- versus *mFus*/*mFUS* mice.

### Homozygous *hFUS* mice display no motor or other overt phenotypes throughout aging, and maintain normal motor neuron counts in the lumbar spinal cord

To provide further insight into the impact of humanisation, *hFUS* mice were aged to 18 months and studied longitudinally to discern if humanising the *FUS* gene had any behavioural phenotypic effect, with particular emphasis on motor function. Humanisation of the FUS gene had no effect on survival of mice heterozygous or homozygous for human *FUS* as shown by a Kaplan-Meier survival curve (**Figure 7A**) and homozygous male and female *hFUS* mice were able to breed successfully (data not shown). SHIRPA was carried out as an observational assessment of *hFUS* mouse phenotype of both sexes at 3- and 18-month timepoints (**Figure S17**). No statistical difference was observed in any of the parameters measured. Body weight measurements show male heterozygous and homozygous *hFUS* mice are marginally, but significantly, heavier than wildtypes, on average by 10% across all timepoints measured. Interestingly this weight difference was not observed in female *hFUS* mice (**Figure 7B**). Neither sex exhibited motor dysfunction as measured by grip strength, nor quantification of leg errors displayed during locotronic analysis (**Figures 7C,D**). Wheel running ability, including the response to a modified wheel presenting a motor challenge, remained equivalent between *hFUS* homozygous and wildtype mice at early and late timepoints. (**Figure 7E**). Analysis of motor neuron numbers in the sciatic motor pool of the lumbar spinal cord showed no differences between 18- month old *hFUS* and wildtype mice (**Figure 7F**). Thus, there was no overt phenotypic differences observed between *hFUS* and wildtype mice, consistent with the very minor transcriptomic impact of *Fus* humanisation.

**Figure 7.**
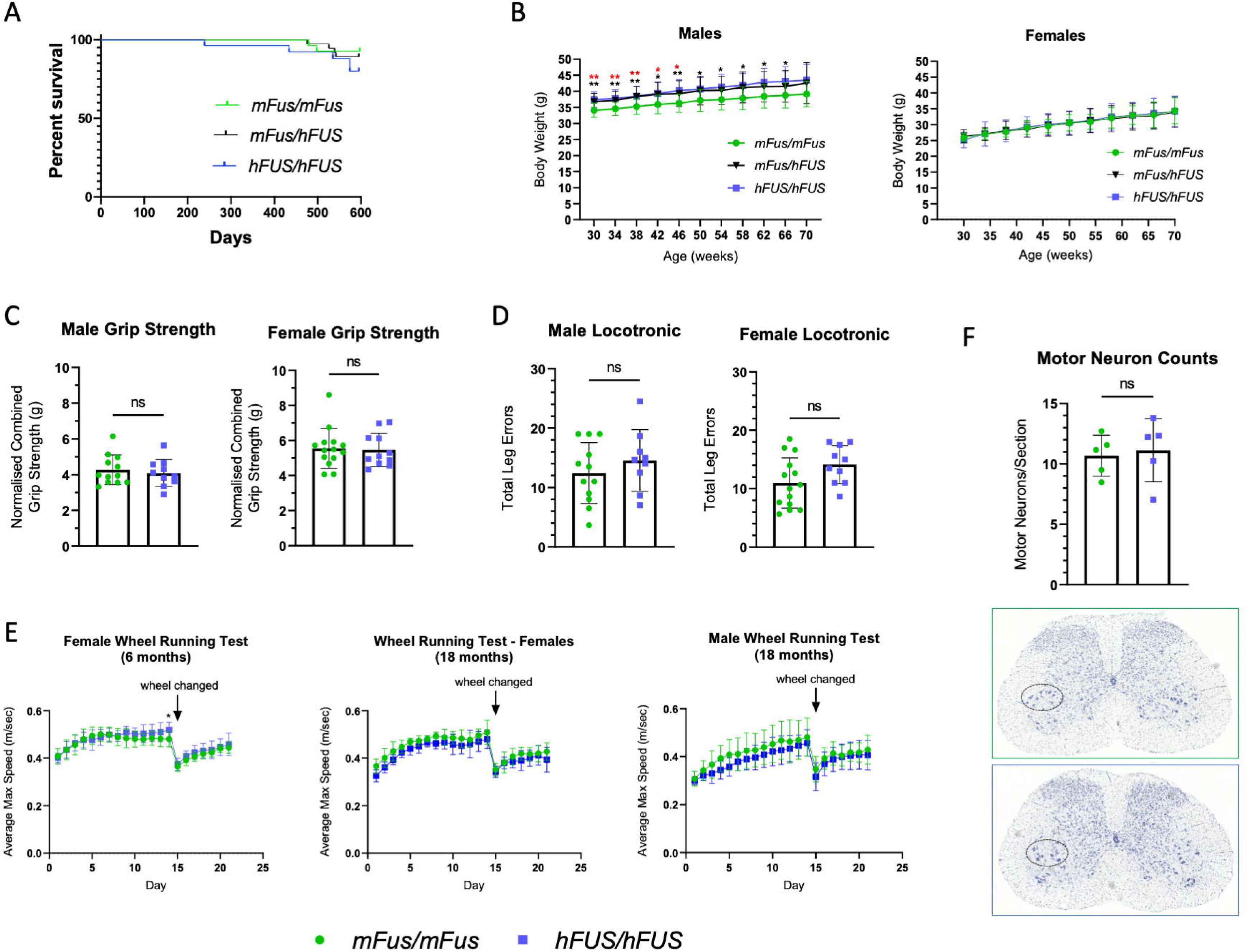
Phenotypic assessments of *hFUS/hFUS* mice show no evidence of an overt phenotype. **(A)** Kaplan-Meier survival curve showing survival data (males and females combined) up to 18 months (n=33, 47, 30; *mFus/mFus, mFus/hFUS, hFUS/hFUS*). **(B)** Body weight measurements of male and female humanized FUS mice. *mFus/mFus* (n=17 male, 16 female), *mFus/hFUS* (n=22 male, 24 female), *hFUS/hFUS* (n=14 male, 13 female), up to 16 months of age. Data presented as mean ± SD, *=p ≤0.05, **=p≤0.01 where black lower* is comparing *mFus/mFus* to *hFUS/hFUS* and red upper* is comparing *mFus/mFus* to *mFus/hFUS,* calculated using a mixed-effects analysis with Dunnett’s post hoc test. **<u>(C)</u>** Grip strength assessment of *hFUS/hFUS* (n=10 male, 11 female) and *mFus/mFus* (n=12 male, 14 female) mice aged 18 months. Data presented as mean ± SD, ns=not statistically significant calculated using Student’s T test. **(D)** Locotronic testing on *hFUS/hFUS* (n=9 male, n=10 female) and *mFus/mFus* (n=12 male, n=14 female) mice aged 18 months. Data presented as mean ± SD, ns=not statistically significant calculated using Student’s T test. **(E)** Motor function via automated wheel running in 6-month old *hFUS/hFUS* (n=13 female) and *mFus/mFus* (n=13 female) mice and 18-month old *hFUS/hFUS* (n=6 female, 8 male) and *mFus/mFus* (n=4 female, 7 male) mice. Data presented as mean ± SD, *=p ≤0.05 calculated using a mixed-effects analysis with Šídák’s post hoc test. **(F)** Motor neuron counts in the ventral horn of the lumbar spinal cord in 18-month old *hFUS/hFUS* and *mFus/mFus* female mice (n=5 per genotype; representative section images shown below). Data presented as mean ± SD, ns=not statistically significant calculated using Student’s T test.

## DISCUSSION

ALS/FTD disorders have no cure or effective treatment and a considerably better understanding of pathomechanisms remains essential for diseases on this spectrum. While cellular and *in vitro* studies are immensely valuable, currently only animal models allow us to look at effects of ageing, systemic effects such as hormones or the microbiome, and environmental interactions (De Giorgio et al., 2019). These studies have also shown, for example, that motor neuron death in ALS is not cell autonomous but entails interaction between neurons and other cell types (Taylor et al., 2016). The range of genome engineering techniques now available enables us to tailor mouse models to the research questions that need to be addressed. To optimise mouse models for ALS/FTD research and the development of new therapies, we describe the replacement of whole coding regions, exons and introns included, of three critically important ALS/FTD genes in mouse (*SOD1*, *TARDBP, FUS*), for their human orthologous sequences, creating ‘genomically humanised’ mice.

We go on to present the essential deep quality control of these new strains, which we regard as a ‘new standard’ in their development as tools for ALS/FTD research, and we provide some phenotype data as proof of principle for the use of these strains.

Few humanised mice exist and there is no ‘best approach’ to how much of a locus should be knocked in; our strategies were bespoke for each gene based on its genomic context and our current knowledge of pathomechanisms. We have created these genomically humanised strains such that the mice express the human splice variants and proteins, noting that on average, human genes express a greater number of splice isoforms than mouse genes (Lee and Rio, 2015), and therefore have increased protein complexity. Furthermore, a small number of amino acid differences within highly conserved mouse and human protein orthologues can significantly affect properties such as propensity to aggregate, for example (Crown et al., 2020; DuVal et al., 2019; Nagano et al., 2015; Perri et al., 2020). Including non-coding gene regions is also important since several known mutations and risk variants exist in such regions (Brown et al., 2021; Dini Modigliani et al., 2014; Ma et al., 2021; Sabatelli et al., 2013). Our genomically humanised strains are ‘knock-in’ animals and therefore express proteins from the endogenous locus at physiological levels. Thus, genomic humanisation delivers mouse models with human protein isoforms and human protein biochemistry, expressed in an endogenous context, with the potential to model mutations and risk variants in coding or non-coding gene regions (Nair et al., 2019; Zhu et al., 2019). Notably, all three lines survive in homozygosity, immediately showing that for the essential genes *FUS* and *TARDBP*, the human genes can functionally replace their mouse counterpart. Aging of hSOD1 is needed to assess functional equivalency of the human gene expressed endogenously in mouse, although we observe no reduction in weight, which would indicate loss of function (data not shown).

Ensuring locus integrity at the nucleotide level is paramount to understanding the impact of precisely engineered alleles in a physiologically relevant context, for which long-read sequencing technology is unparalleled for the task. We demonstrate that long-read sequencing technologies and analysis tools are readily accessible, setting a new standard for large-engineered allele quality control in an age of expanding genome engineering capabilities.

In addition to confirming the integrity of our precisely engineered alleles, we took advantage of the strain and sequence differences between the targeting vector and ES cells to map homologous recombination breakpoints. The full length of the long homology arms used, tens of kb in length, were frequently not fully integrated into the locus together with the human gene, suggesting that future strategies may be successful with far shorter regions of homology. Interestingly, we note that for *Tardbp,* breakpoints are closest to the humanised region (11kb 5’ proximal and 5.5kb 3’ distal) and this was the only strain for which we had used CRISPR/Cas9 assisted homology directed repair, with guides cutting specifically within the mouse endogenous *Tardbp* loci. In all strains, we find that recombination breakpoints are free from indels or any other type of mutation.

Looking to the effect on humanised gene products, h*SOD1* expression is not changed in the CNS, but reduced at the mRNA and protein levels only in skeletal muscle. As CNS expression levels are remarkably similar between wildtype and *hSOD1*, this argues against general disruption caused by the conditional element of the allele. One possible explanation for the reduction in skeletal muscle *SOD1* expression could be the presence of tissue specific transcriptional control elements in the humanised non-coding regions (introns and/or 3’ UTR). Regardless of the mechanism, this finding clearly warrants further investigation. For h*TARDBP*, the minor mRNA expression changes detected did not translate into differences at the protein level or in endogenous TDP-43 splicing function, suggesting that human TDP-43 can functionally replace the mouse protein. Finally, for h*FUS* mice, the moderate reductions in *FUS* mRNA had little consequence at the genome-wide transcriptomic level or phenotypic level, functionally corroborating that expression levels detected lie within a physiological range.

These changes underlie the need for characterisation of the humanised lines before ALS/FTD mutations are introduced, in order to set a baseline phenotype and provide appropriate humanised-non-mutant controls, whilst further investigation could lead to insights regarding human-specific gene regulation.

To gain further insight into the impact of humanisation, we have more comprehensively characterised *hFUS* mice at the behavioural and molecular level. hFUS homozygotes are fully viable, fertile, and survive in equal measure to wildtype mice through to 18 months of age. We found no motor phenotypes, nor loss of motor neurons in the lumbar spinal cord, in contrast to reports of transgenic hFUS expressing mice (also without mutations) that show motor phenotypes and motor neuron loss (Ling et al., 2019; Mitchell et al., 2013). While we cannot rule out neurodegenerative phenotypes developing beyond this timeframe, it nevertheless shows remarkable functional synergy with mouse *Fus,* and provides a robust foundation for the introduction of pathogenic mutations for future study. Looking at body weight data, heterozygous and homozygous *hFUS* males have marginally, but consistently larger body mass than wildtype, which may be related to subtle differences in gene expression levels, but ultimately is a minimal phenotypic difference. Taking phenotypic, transcriptomic, and expression data together, clearly these lines are not 100% identical to wildtype animals, which underscores the need to use them as the most accurate genetic control for future mutant lines, in order to assess the impact of mutations specifically, distinct from the impact of humanisation. Further characterisation of the h*SOD1* and h*TARDBP* mice at the behavioural and molecular level throughout their entire lifespan will be required to complete their characterisation.

These three strains are the first KI lines in which the mouse expresses the human SOD1, TDP-43, and FUS protein from the endogenous locus. These strains will be of great utility for understanding the biology and dysfunction of these proteins when mutated, and is particularly important for the highly dosage-sensitive genes *FUS* and *TARDBP*. The *hTARDBP* strain is the first model to express only human protein without endogenous mouse protein as, to our knowledge, all attempts to rescue the KO phenotype with transgenics (including BACs) have been unsuccessful.

Furthermore, these humanised models will be an important resource for the validation of ASOs, the highly promising therapeutics aimed at modulating human gene expression levels, as these require the exact human sequence for preclinical trials. Moreover, the humanised mice will uniquely allow us to test the possible effects of ASOs, or any other therapeutic agents, on systemic long-term loss of function that cannot be assessed in transgenic models. This will be critically important as all three genes are ubiquitously expressed and perform crucial biological functions; recent failed clinical trials from ASOs aimed against lowering Huntingtin (*HTT*) showed adverse effects (Kwon, 2021) that might be mediated by *HTT* loss of function, highlighting the growing need for improved pre-clinical understanding of the risks associated with gene therapies long-term (Editorial, 2021).

### Limitations

We need a variety of models to study the complexities of ALS and FTD. Despite the limitations of the mouse as a system to model human neurodegenerative disease (including differences in axon length; rate of ageing; synaptic connections between upper and lower motoneurons; anatomy of neuromuscular junctions), mice will continue to be invaluable to study fundamental disease processes because they provide a complex *in vivo* environment that is critical for the study of interactions between multiple tissues and cell types (motoneurons, glia, muscle and other non-neuronal tissues) that are likely necessary for disease development and progression. By generating these humanised mice for ALS/FTD research, we aim to produce, at least at the biochemical level, the closest models to the human condition to study the impact of human genes and proteins within a complex mammalian organism. These new humanised models are not designed to entirely replace existing models. Physiological models for ALS/FTD (i.e. those that involve study of genes at endogenous loci), typically have slow onset of disease and may not reach end-stage; therefore, to address questions surrounding late or end-stage disease, transgenic overexpression models may be better suited.

In general, a potential critical limitation of genomically humanised mice is that replacing the endogenous mouse gene by its human *wildtype* counterpart could lead to functional consequences. As a proof of principle, we present an extensive characterisation of h*FUS* mice showing that although minimal, some differences exist between h*FUS* mice and littermate controls. Taking phenotypic, transcriptomic, and expression data together, clearly these lines are not identical to wildtype animals, which underscores the need to use them as the most accurate genetic control for future *mutant* lines, in order to assess the impact of mutations specifically, distinct from the impact of humanisation. Further characterisation of the h*SOD1* and h*TARDBP* mice at the behavioural and molecular level throughout their entire lifespan will also be required to complete their characterisation. Moreover, humanised mice (as with other mutant mouse strains) will require continued quality control to make sure that the human allele remains intact through the process of colony maintenance.

With the three humanised lines, we decided to maintain the mouse promotor and upstream non-coding exons. The rationale for this approach was to minimize disruption of expression from the endogenous locus within the context of the mouse. Thus, these models will not allow us to study the possible effects on expression of the human promoters regions (or variants within), although we note further modifications can be made to the wildtype humanised animals, which are essentially templates for further changes.

Looking to the future, single humanised alleles may not be sufficient to study disease mechanisms involving binding partners of humanised proteins, or to assess human specific splicing events (Brown et al., 2021; Ma et al., 2021), and new bespoke humanised alleles may need to be engineered and cross-bred for these purposes. In turn these may aid in more faithfully recapitulating later stage pathology and phenotypes, for further mechanistic insight in vivo and for improved read outs for testing therapeutic strategies.

Looking to the future, single humanised alleles may not be sufficient for digging deeper into mechanisms involving binding partners of humanised proteins, and new bespoke humanised alleles may need to be engineered and cross-bred for these purposes. In turn these may aid in more faithfully recapitulating later stage pathology and phenotypes, for further mechanistic insight in vivo and for improved read outs for testing therapeutic strategies. The three humanised alleles presented here are an important first step and proof of principle that human ALS/FTD genes can functionally replace their mouse orthologues enabling the introduction of mutations to better understand the impact of human mutant or variant genes and proteins.

### Conclusions

Here, we present the first genomically humanised KI mouse models for three ALS/FTD genes, *SOD1*, *TARDBP*, *FUS*. We demonstrate the utility of a pipeline for the robust characterisation of complex KI alleles consisting of indirect capture for enrichment of the region of interest followed by long-range sequencing. The strategy provides confidence in the integrity of the allele at the sequence level, confirms that gene targeting occurred at the correct locus for all three lines, including mapping of the homologous recombination events. In all cases, the human gene functionally replaced the mouse endogenous gene. Moreover, human *FUS* can functionally replace the mouse gene throughout the ageing process. We envisage that these mice, which will be freely available from the European Mutant Mouse Archive (EMMA), may be mutated by CRISPR or other approaches to recreate human ALS/FTD causal mutations, to dissect pathogenesis including aggregation, and for developing new therapies, and we encourage their uptake by the community.

## MATERIALS AND METHODS

### Mice

#### Animal housing and maintenance

All mice were maintained and studied according to UK Home Office legislation at MRC Harwell (Home office licence 20/0005). Mice were fed *ad libitum* (Rat and Mouse Breeding 3 (RM3), Special Diet Services) with free access to water (chlorinated to 9-13 ppm). Mice were kept under constant conditions with a regular 12 h light (19:00-07:00) and dark (07:00-19:00) cycle, temperature of 21±2°C and humidity 55±10%. Mice were housed in same sex cages of up to five mice. Cages contained Aspen Chips bedding (Datesand), shredded paper and a rodent tunnel for enrichment. All three humanised mice strains are maintained by backcrossing humanised heterozygous animals to C57BL/6J mice. For all the experimental cohorts presented, mice were obtained by intercrossing heterozygous humanised animals to obtain all three genotypes within a single cross, with randomised weaning into cages.

### Genomic humanisation and strain names

#### Generation of genomically humanised *SOD1* mice

The official allele designation is *Sod1* ^tm1.1(SOD1)Emcf^ (MGI:6403195), referred to here as *hSOD1*. A BAC targeting construct harbouring the mouse *Sod1* locus (BAC B6Ng01-068O19; strain C57BL/6N) was engineered to replace the mouse *Sod1* gene from the ATG start site (g.117) to the end of the 3’UTR (g.5583) with the orthologous human *SOD1* genomic sequence from a BAC harbouring the human *SOD1* locus (BAC RP11-535E10). *LoxP* sites were inserted into intron 3, and 1kb downstream of the inserted human 3’UTR (**Figure 1A**). The sequence between the *loxP* sites was duplicated to allow conditional expression of a second copy of the sequence. An *FRT*-flanked Neomycin cassette was inserted between the first and second copies of the duplicated sequence. This construct was electroporated into the 129X1/SvJ-129S1/SV mouse ES cell line R1, humanising *Sod1* in the mouse genome via homologous recombination. Correctly targeted clones were initially identified through Loss-Of-Allele (LOA) copy via qPCR and ddPCR assays, while karyotype of positive clones was then screened employing ddPCR (Codner et al., 2016; Valenzuela et al., 2003). Mice were generated by injection of modified ES Cells into donor blastocysts, with the resultant chimeric male offspring crossed to C57BL/6J females to obtain germline transmission (GLT). Following confirmation of GLT, hSOD1 heterozygous mice were backcrossed for one further generation to the C57BL/6J strain. Selection cassette removal was performed at generation N3 C57BL/6J backcrossed animals, via cytoplasmic injection of IVF derived 1-cell embryos with *Flpo* mRNA. The Neo-negative line was then backcrossed for one further generation onto C57BL/6J, followed by intercrossing to produce animals at generation N4 for long-read sequencing and preliminary expression analysis. Genotyping was performed by LOA copy number qPCR using custom probe sets: Mouse *Sod1 allele,* forward primer GTACCAGTGCAGGACCTCAT, reverse primer AGCGTGCTGCTCACCTCT, 5’-FAM probe AACATGGTGGCCCGGCGGATG; hSOD1 allele, forward primer GTGCAGGTCCTCACTTTAATCC, reverse primer CCAGAAAGCTATCGCCATTATTACAAG, 5’-FAM probe CCAAAGGATGAAGAGAGGTAACAAGATGC. This line has been submitted to the European Mouse Mutant Archive (EMMA, EM:13075).

#### Generation of genomically humanised *TARDBP* mice

The official allele designation is Tardbp^em2.1(TARDBP)H^ (MGI:6513996), referred to here as hTARDBP. A B AC targeting construct harbouring the mouse *Tardbp* locus (BAC B6Ng01-103M13) was engineered to replace the mouse *Tardbp* gene from the ATG start site to the TAG stop codon with the orthologous human *TARDBP* genomic sequence from a BAC harbouring the human *TARDBP* locus (BAC RP11-829B14; strain C57BL/6N); an *FRT*-flanked Neomycin cassette was inserted into intron 3 of the gene (**Figure 2A**). This construct was electroporated into the 129X1/SvJ-129S1/SV mouse ES cell line R1, humanising *Tardbp* in the mouse genome via homologous recombination assisted by CRISPR-Cas9 (delivered by co-electroporation with px459 plasmids; sgRNA guide insert sequences: gCTCCACCCATATTACCACC and gGTCGGGCCCATCTGGGAATA). Correctly targeted clones were initially identified through Loss-Of-Allele (LOA) copy via qPCR and ddPCR assays, while karyotype of positive clones was then screened employing ddPCR (Codner et al., 2016; Valenzuela et al., 2003). Mice were generated by injection of modified ES Cells into donor blastocysts, with the resultant chimeric male offspring crossed to C57BL/6J females to obtain GLT. Following confirmation of GLT, *hTARDBP* heterozygous mice were bred one further generation to the C57BL/6J strain, followed by intercrossing to generate homozygotes for long-read sequencing and preliminary expression analysis. Cohorts used for analysis here still contain the selection cassette. Genotyping was performed by Loss-Of-Allele (LOA) copy number qPCR using the commercially available TaqMan probesets Mm00139953 (mouse), Mm00735064 (mouse), Hs06599429 (human), Hs06560655 (human) (ThermoFisher).

#### Generation of genomically humanised FUS mice

The official allele designation is Fus^tm3.1(FUS)Emcf^ (MGI:6193752), referred to here as hFUS. A BAC targeting construct harbouring the mouse *Fus* locus (BAC RP24-297F14; strain C57BL/6J) was engineered to replace the mouse *Fus* gene from the ATG start site (g.106) to the end of the 3’UTR (g.14574) with the orthologous human *FUS* genomic sequence, and an *FRT*-flanked Neo cassette was inserted 1kb downstream from the humanised 3’UTR (**Figure 3A**). This construct was electroporated into the 129X1/SvJ-129S1/SV mouse ES cell line R1, humanising *Fus* in the mouse genome via homologous recombination. Correctly targeted clones were initially identified through Loss-Of-Allele (LOA) copy via qPCR and ddPCR assays, while karyotype of positive clones was then screened employing ddPCR (Codner et al., 2016; Valenzuela et al., 2003). Mice were generated by injection of modified ES cells into donor blastocysts, with the resultant chimeric male offspring crossed to C57BL/6J females to obtain GLT. Following confirmation of GLT, *hFUS* heterozygous mice were bred one further generation to the C57BL/6J strain, followed by crossing to the Gt(ROSA)26Sortm2(CAG-flpo,-EYFP)Ics line (C57BL/6Ntac background) to excise the *FRT*-flanked Neo selection cassette. The Neo-negative line was then backcrossed for at least six further generations onto the C57BL6/J background before homozygotes were generated for long-read sequencing and experimental cohorts were bred. Genotyping was performed by LOA copy number qPCR using the commercially available TaqMan probesets Mm00217486_cn (mouse), Mm00217500_cn (mouse), Hs02670898_cn (human), Hs05441121_cn (human) (ThermoFisher). This line has been submitted to the European Mouse Mutant Archive (EMMA, EM:13073).

### Allele verification pipeline

#### Xdrop enrichment, Oxford Nanopore sequencing and analysis

Xdrop^TM^ enrichment, amplification, and Oxford Nanopore sequencing was performed by Samplix Services (Denmark) (Blondal et al., 2021). High molecular weight DNA was extracted from homozygous humanised mouse tissue using SDS/proteinase K lysis buffer (100 mM NaCl, 10 mM Tris-Cl pH 8, 25 mM ETDA pH 8, 0.5% SDS, 20 µg/ml RNAse A, 100 µg/ml proteinase K) and phenol/chloroform purification (gentle mixing/inversion only, no vortexing), followed ethanol precipitation and elution in 10 mM Tris-Cl pH 8.5. DNA was evaluated by Tapestation 2200 System (Agilent Technologies Inc.), using Genomic DNA ScreenTape according to the manufacturer’s instructions (average sizes: *hSOD1* >100 kb, *hTARDBP* >30 kb, *hFUS* >60 kb). A series of ‘Detection Sequence’ droplet PCR (dPCR) assays, and accompanying qPCR validation assays in the adjoining region, were designed to span each locus (**Table S1; Figures S2,S3,S4**). Primer efficiency of the assays was tested by serial sample dilution in Samplix Primer test PCR kit according to manufacturer’s instructions. 8-9 ng DNA was partitioned in droplets by Xdrop™ and subjected to dPCR using the Detection Sequence assay. The droplet productions were then stained and sorted by fluorescence-activated cell sorting (FACS). DNA was then released from the isolated droplets, and DNA was again partitioned in droplets by Xdrop™ and amplified by droplet multiple displacement amplification (dMDA) as previously described (Blondal et al., 2021). After amplification, DNA was isolated and quantified, and enrichment was validated by qPCR (data not shown) before Oxford Nanopore Sequencing.

Minion Oxford Nanopore Sequencing libraries were prepared from the dMDA samples as described by the manufacturer’s instructions for Premium whole genome amplification protocol (SQK-LSK109) with the Native Barcoding Expansion 1-12 (EXP-NBD104) including a T7 endonuclease I digestion step for debranching followed by bead purification (MagBio). Generated raw data (FAST5) was subjected to base-calling using Guppy v.3.4.5 with high accuracy and quality filtering to generate FASTQ sequencing data (Samplix Services). 3-5 Gb of sequencing data was obtained for each sample. Subsequently, the data was aligned with minimap2 (Li et al. 2018) to the mouse reference C57BL/6J genome (GRCm38.p6), the mouse reference strain 129 genome (129S1_SvImJ) the human reference genome (GRCh38.p13), and custom reference sequences representing the intended allele sequences. In addition, the *hTARDBP* sample was aligned using NGMLR (Sedlazeck et al., 2018). Samtools was used to convert, sort, and index alignment data files for visualization in IGV (Robinson et al. 2011). Bioinformatics commands for the alignments can be found in **Figure S18**.

### Gene expression

#### Tissue isolation

Mice were culled via intraperitoneal injection of pentobarbitone and extracted tissues were stored at −80°C until processed. Lumbar spinal cord tissue for staining was collected by hydraulic extrusion of the whole spinal cord into cold PBS. The lumbar region was identified visually, cut, and embedded in OCT and frozen over isopentane and dry ice.

#### RT-PCR and Quantitative real-time PCR

Total RNA was isolated from frozen tissue using RNeasy lipid or fibrous tissue mini kits (Qiagen) or Trizol reagent (ThermoFisher) following manufacturer’s instructions. cDNA was synthesised using MultiScribe Reverse Transcriptase kit (ThermoFisher) or qScript cDNA Synthesis kit (QuantaBio) following manufacturer’s instructions. For RT-PCR, human- and mouse-specific primer pairs were designed for each gene: *hFUS* forward (5’-CGGTCGTCTGGAACTTTGTT-3’) and reverse (5’-CCATAACCGCCACCACTG-3’); mouse *Fus* forward (5’-CGGTCGTCTGGAACTTTGTT-3’) and reverse (5’-CATAACCACCACCGCCAC-3’); *hSOD1* forward (5’-TCGTCTTGCTCTCTCTGGTC-3’) and reverse (5’-CAGGCCTTCAGTCAGTCCTT-3’); mouse *Sod1* forward (5’-AACCAGTTGTGTTGTCAGGAC-3’) and reverse (5’-CCACCATGTTTCTTAGAGTGAGG-3’); *hTARDBP* forward (5’-ATGACTGAGGATGAGCTGCG-3’) and reverse (5’-CACAAAGAGACTGCGCAATCTG-3’); mouse *Tardbp* forward (5’-CCATTCAGAGCTTTTGCCTTC-3’) and reverse (5’-AGCTCCACCCCCTCTACTG-3’). PCR reaction products were run on 1.5% agarose gels to assess for the presence/absence of mouse or human gene products.

To quantitatively analyse humanised gene mRNA expression, plus *Masp2* mRNA, qRT-PCR was performed using Fast SYBR Green Master Mix (ThermoFisher) or MasterMix qPCR Lo-ROX (PCR-Bio) and 200 nM or 250 nM of each forward and reverse primer. The thermal amplification conditions were: 95°C for 20 s, then 40 cycles of 95°C for 3 s and 60°C for 30 s. The specificity of primer binding and amplification was confirmed by melt curve analysis. To control for non-specific amplification, no-template reactions were performed using all reagents except the sample. qRT-PCR primer pairs for *Sod1*, *Tardbp,* and *Fus* were 100% conserved between mouse and human: h*FUS/Fus* forward (5’-CACAGGCTATGGAACTCAGTCA-3’) and reverse (5’-TGAAGAGAGGCATGTTGGAGA-3’); h*SOD1/Sod1* forward (5’-CACAGGCTATGGAACTCAGTCA-3’) and reverse (5’-CCACCTTTGCCCAAGTCATC-3’); h*TARDBP/Tardbp* forward (5’-GAGCCTTTGAGAAGCAGAAA-3’) and reverse (5’-CCACCTGGATTACCACCAAA-3’). qRT-PCR primers were also used to analyse *Masp2* expression: *Masp2* forward (5’-ACCGCTGCGAGTATGACTTT −3’) and reverse (5’-CCTGTGAACGGCTTCTCATT −3’). *S16* or *GAPDH* expression were used as internal controls: *S16* forward (5’-TTCTGGGCAAGGAGCGATT-3’) and reverse (5’-GATGGACTGTCGGATGGCA-3’); *GAPDH* forward (5’-CGGCCGCATCTTCTTGTG-3’) and reverse (5’-CCGACCTTCACCATTTTGTCTAC-3’). qRT-PCR reactions were carried out on 7500, QuantStudio or Roche Lightcycler 480 II Real-Time PCR machines. qRT-PCR data was analysed using the manufacturer’s analysis software following the 2^-ΔCt^ method.

#### Protein expression analysis

For FUS and SOD1, tissue was disrupted in Pierce RIPA buffer (ThermoFisher) + protease inhibitors (Roche/Sigma), whereas TDP-43 was disrupted in Urea 7M-1% SDS, followed by matrix D lysing tubes for 2x 30 s at 5500 rpm (Precellys). Homogenates were centrifuged at 13,000x g for 10 minutes at 4°C. Supernatant (total protein) was aliquoted and stored at −80°C. Genotypes studied included wildtype, heterozygous and homozygous mice; at least n=3 per genotype for different tissues, including: brain, spinal cord, skeletal muscle and liver; except for spinal cord and TA for *TARDBP* humanised mice where n=2 was used. Protein concentration was assessed using a DC assay (BioRad) or BCA assay (Sigma). FUS protein extracts were ran on 4-12% Bis-Tris protein gels (NuPAGE, ThermoFisher) with 1X MOPS (ThermoFisher) + 500 μ antioxidant (ThermoFisher) at 200V. SOD1 protein extracts were ran on Novex 14% Tris-Glycine protein gels (ThermoFisher) with 1X Tris-Glycine SDS (ThermoFisher) at 200V. FUS and SOD1 protein was transferred to a nitrocellulose membrane using iBlot Transfer Stacks and iBlot machine. For TDP-43, protein extracts were run on 10% Bis-Tris gels with Tris-Glycine SDS running buffer at 150V, followed by wet transfer with running buffer and 20% methanol at 280 mA. For SOD1, membranes were reversibly stained for total protein using a Revert 700 Total Protein Stain Kit (LI-COR) following the manufacturer’s instructions. After incubation with 4% skimmed milk (Sigma Aldrich) in PBST (0.1% Tween-20) for 1.5 h, membranes were incubated with the following antibodies: FUS 562 (a rabbit polyclonal raised against a peptide matching the C-terminus of both mouse and human FUS; Novus NB100-562, 1:10,000), pan TDP-43 (a mouse monoclonal antibody recognising an epitope of the N-terminus of TDP-43 that is identical between mouse and human, Bio techne, MAB7778, 1:1000), human specific TDP43 (a mouse monoclonal against the C-terminus of TDP-43 that recognises human TDP-43 protein but not mouse TDP-43, Proteintech 60019-2-Ig, 1:1000), pan SOD1 (a custom-made goat polyclonal antibody recognising a C-terminal epitope of SOD1 that is identical between mouse and human proteins; 1:1000; kind gift from Dr Jonathan Gilthorpe, Umea University, Sweden), and GAPDH (ThermoFisher AM4300, 1:2000 or Proteintech 60004-1-Ig, 1:1000) diluted in 4% milk/PBST either O/N at 4°C or for 1.5 h at RT. Membranes were washed 3x 5 minutes in PBST then incubated with anti-rabbit and anti-mouse antibodies (LI-COR, 1:15,000 or ThermoFisher, 1:10,000) diluted in 4% milk/PBST for 1.5 h at RT. Membranes were washed 2x 5 minutes in PBST and a further 5 minutes in PBS before being dried (10 minutes) and imaged on a LI-COR Odyssey Scanner or an Image Quant LAS4000, GE healthcare. Blots were quantified using Empiria Studio 1.3 (LI-COR) or image count TL imaging software (GE healthcare). Blot images from *hFUS* samples are shown in **Figure S16**.

#### Immunohistochemistry of motor cortex and image analysis

*hFUS* brain (8-months) was dissected into left and right sides and again into anterior and posterior portions, drop fixed in 4% PFA for 24 hours, washed in PBS 3 x 10 minutes, equilibrated in 30% sucrose, and cryopreserved in OCT over isopentane on dry ice. Serial transverse sections (20 µm) were cut through the motor cortex region of the anterior embedded brain tissue on a cryostat, and further fixed in PFA for 10 minutes. Sections were washed 3x 10 minutes in PBS then incubating in blocking buffer (10% normal goat serum in TBST, 0.2% Triton) for 1 h. Sections were washed 3x 10 minutes in PBS then incubated with an anti-FUS 565 antibody (NB100-565; 1:500, Novus) in blocking buffer overnight at 4°C. Sections were washed 3x 10 minutes in PBS then incubated with anti-mouse IgG (1:1500, AlexaFluor 488, ThermoFisher) in blocking buffer for 2 h. Sections were then washed 3x 10 minutes in PBS and mounted using Prolong Glass Antifade Mountant with NucBlue for nuclear counterstaining (ThermoFisher). Snap images were obtained using an inverted confocal microscope (Zeiss).

### RNA sequencing

Whole spinal cords were extracted via hydraulic extrusion of 14-week old male animals and stored in RNAlater (ThermoFisher); genotypes included wildtype, *hFUS* homozygotes, and *Fus* KO heterozygotes; n=4 per genotype. Total RNA extraction, library preparation, Illumina sequencing, and bioinformatics analysis was performed by Lexogen services. Total RNA extraction using Lexogen SPLIT kit; Library preparation using CORALL Total RNA-Seq Library Prep Kit with Poly(A) selection; SR75 High Output sequencing on Illumina NextSeq 500. Data was analysed using the Lexogen CORALL Data Analysis Pipeline, including sequence QC (fastqc), read trimming (cutadapt), mapping, quantification, and differential expression. FASTQ files of raw reads were mapped to the GRCm38 mouse genome using STAR. Gene and transcript expression were estimated with Lexogen’s Mix2 software. Transcript level differential expression was performed using DESeq2.

### SNP analysis

Transnetyx services performed MiniMUGA SNP analysis; an array-based platform with over 10,000 SNP markers that can be used to determine genetic background in 241 inbred strains of mice.

### Phenotyping

Phenotyping tests on *hFUS* mice were carried out at the same time each day and mice were acclimatised for 30 minutes prior to each test. The same experimenter carried out each test and was blind to genotypes of the mice.

#### SHIRPA

Modified SHIRPA (Rogers et al., 1997) was carried out at 3 months and 18 months; 3 months: n=17,15,16,13 male *mFus/mFus,* male *hFUS/hFUS,* female *mFus/mFus,* female *hFUS/hFUS,* 18 months: n=12,10,14,11 male *mFus/mFus,* male *hFUS/hFUS,* female *mFus/mFus,* female *hFUS/hFUS*). Any significant differences observed between genotypes for SHIRPA parameters were assessed using Fisher’s exact test and chi-square tests.

#### Weight checks

Mice were weighed inside a beaker, without the use of anaesthetic, every four weeks and before relevant *in vivo* tests.

#### Grip strength

Grip strength of 22 males (n=12 wildtype and n=10 *hFUS* homozygous mice) and 25 females (n=14 wildtype and n=11 *hFUS* homozygous mice) aged 18 months was assessed using the force sensor equipment as per manufacturer’s instructions (Bioseb Grip Strength Meter). Mice were lowered onto the grid so only the forelimbs could grip, and were pulled steadily downward. This was repeated 3 times. Mice were lowered onto the grid allowing fore- and hind limbs to grip, and were pulled steadily downward. This was repeated 3 times.

#### Locotronic

Locotronic foot misplacement analysis of 22 males (n=12 wildtype and n=10 *hFUS* homozygous mice) and 25 females (n=14 wildtype and n=11 *hFUS* homozygous mice) aged 18 months was assessed as per manufacturer’s instructions (Intellibio). The equipment contains a horizontal ladder within a corridor and mice are motivated to move from the illuminated start area to the darker finish area. Infrared sensors above and below the ladder detect errors in paw placement. Three runs were carried out per mouse with at least 30 minutes between runs, and the mean average total leg errors calculated. Runs which took longer than 30 s after exiting the start area were discounted (Stewart et al., 2019).

#### Wheel-running motor function (MO)

Motor function of 26 female mice (n=13 wildtype and n=13 *hFUS* homozygous mice) aged 6 months and 18 males (n=9 wildtype and n=9 *hFUS* homozygous mice) and 18 females (n=9 wildtype and n=9 hFUS homozygous mice) aged 18 months was assessed using an automated wheel-running test (Mandillo et al., 2014). Mice were singly housed in cages containing a voluntary running wheel attached to a computer for recording data for three weeks. Food and water were available *ad libitum*. After two weeks, the wheel was changed for one with rungs missing at irregular intervals, presenting a motor challenge. Only data recorded during the dark period (19:00-07:00) was analysed. Average maximum wheel running speed was calculated (in 5-minute bins) and analysed. Any mice which did not complete the three weeks were removed from analysis; in particular, aged mice are more prone to weight loss with this test, which is cause for removal before they reach humane endpoints.

#### Motor neuron counts in lumbar spinal cord

5 female *mFus/mFus*, and 5 female *hFUS/hFUS* mice were sacrificed at 18 months, spinal cord was removed via hydraulic extrusion, and a 1 cm section of the lumbar spinal cord bulge (centring on the widest point of the lumbar enlargement) was dissected and embedded in OCT and frozen over isopentane on dry ice. Serial transverse sections (20 μm) were cut from regions L1 to L6 of the lumbar spinal cord and collected onto glass slides. Every third section was analysed leaving a gap of 60 μm between sections and ensuring the same motor neuron was not counted twice. Slides were stained for 20 minutes in Gallocyanin (0.3 g gallyocyanin, 10 g chrome alum, distilled water up to 100 ml), rinsed with water, dehydrated and mounted using CV Ultra Mounting Media (Leica Biosystems). Sections were scanned using a Nanozoomer slide scanner (Hamamatsu) and motor neurons counted. Motor neurons with the following criteria were counted: a dense, visible nucleolus, a diameter of >15 μ and visible dendritic branching. 35-40 sections were counted per animal and the level of the spinal cord standardised by morphological assessment, such that equivalent sections from L1-L6 were included for counting, centring on L4-L5.

### Statistical analysis

Statistical analysis was conducted using GraphPad Prism and SPSS. Refer to figure legends for individual tests used.

## DECLARATIONS

### Ethics approval and consent to participate

Animal procedures performed at the MRC Harwell Institute were licenced by the Home Office under the Animals (Scientific Procedures) Act 1986, UK and additionally approved by the relevant Institutional Ethical Review Committees.

### Consent for publication

“Not applicable”.

### Availability of data and materials

The datasets used and/or analysed during the current study are available from the corresponding author on reasonable request.

### Competing interests

The authors declare that they have no competing interests.

### Funding

TJC, RRN, CT, RM, SG, EMCF were funded by the UK Medical Research Council (MRC) (MC_EX_MR/N501931/1). GP, DT, ZA were funded by MRC PhD studentships. AA, GFC, EO, JH, GA, MS, and LT were funded by the UK Medical Research Council (MRC, award A410). FdG was funded by a Motor Neurone Disease Association PhD studentship. RB-S and AD were funded by the UK MRC (MR/L021056/1) and the UK Motor Neurone Disease Association. AA-A was funded by the ISCiii Miguel Servet programme, by the ISCiii (PI17/00144 and PI20/00422), and CIBERNED. EMCF was funded by the Rosetrees Trust.

### Authors’ contributions

Conceptualization (EMCF, AA-A, AD, TJC), Supervision (EMCF, TJC, AA-A, LT), Construct Engineering and QC (AD, FdG, RB-S, MS), Gene Targeting and Mouse Generation (LT, GFC, AA), Colony Supervision (JH, GA, MS, GP, DT, TJC), Long-Read Sequencing Analysis (TJC, DT, EON), Gene Expression Studies (GP, DT, JN, JMB-A, AM-G, SG, TJC, SC, ZA), Behavioural Phenotyping (GP, SG, RRN, CT, DT, JH, GA, MS, TJC), Motor Neuron Counts (GP), RNAseq analysis (TJC, GP), SNP Analysis and Sanger sequencing (GP, JN, RM, TJC), Manuscript Writing (TJC, EMCF, AA-A, GP, DT), Acquired Funding (EMCF, AA-A). All authors read and reviewed the manuscript.

## Acknowledgements

We thank all staff involved at the Mary Lyon Centre, MRC Harwell Institute, for animal generation, assistance with phenotyping, and staff on ward 5 for animal husbandry. We thank the histology core at MRC Harwell institute for contributions to motor neuron counting. We thank Dr Bernadett Kalmar and Professor Linda Greensmith for advice and direction for motor neuron counting. We thank Dr Cecilie Nyholm Andersen, Dr Ronni Schmidt Bertelsen, and Samplix Services, Denmark, for Xdrop genomic DNA target capture, Nanopore sequencing and initial processing of raw data, and assistance with long-read sequence alignments. We thank Professor Pietro Fratta for critical reading of the manuscript.

---

**Figure S1. Brief overview of Xdrop locus capture.** Using the *hFUS* locus as an example. High molecular weight DNA was encapsulated in Xdrop droplets; detection sequences (indicated in green) were designed to be spaced such that the size of the captured DNA would allow overlapping coverage; droplet PCR and staining followed by FACS isolated droplets from the target detection loci; droplet multiple displacement amplification to amplify target DNA; followed by DNA purification and Oxford Nanopore sequencing.

**Figure S2. Alignment of *hSOD1/hSOD1* sequencing reads to 1 Mb *Sod1* C57BL/6J locus.** IGV visualization of minimap2 alignment to the GRCm38 C57BL/6J reference genome, zoomed in to 1 Mb surrounding the *Sod1* locus. Red asterisks note the positions of detection sequences, which broadly correspond to the peaks in coverage.

**Figure S3. Alignment of *hTARDBP/hTARDBP* sequencing reads to 1 Mb *Tardbp* C57BL/6J locus.** IGV visualization of minimap2 alignment to the GRCm38 C57BL/6J reference genome, zoomed in to 1 Mb surrounding the *Tardbp* locus. Red asterisks note the positions of detection sequences, which broadly correspond to the peaks in coverage.

**Figure S4. Alignment of *hFUS/hFUS* sequencing reads to 1 Mb *Fus* C57BL/6J locus.** IGV visualization of minimap2 alignment to the GRCm38 C57BL/6J reference genome, zoomed in to 1 Mb surrounding the *Fus* locus. Red asterisks note the positions of detection sequences, which broadly correspond to the peaks in coverage.

**Figure S5. *hSOD1* allele theoretical map alignment.** Alignment (minimap2) of *hSOD1/hSOD1* mouse sequencing reads to a file of the intended humanised allele sequence (green outline top IGV panel); a single imperfect alignment feature (*h) maps the correct base in the majority, but is within a homopolymeric sequence and shows a higher error rate within reads. Engineered duplications (red and brown arrows) and engineered *loxP* and *FRT* site insertions are only evident as anomalies when aligning to human and mouse reference genomes (lower IGV panels). In the alignment to the human genome, a SNP is evident in intron 1 (#) that was known to be present in the targeting construct and is a known human polymorphism.

**Figure S6. *hTARDBP* allele theoretical map alignment.** Alignment (minimap2) of *hTARDBP/hTARDBP* mouse sequencing reads to a file of the intended humanised allele sequence, including the *FRT*-flanked selection cassette, which was not excised before sequencing (green outline top IGV panel). The selection cassette region is only evident as an anomaly when aligning to human reference genome (*sc; human reference genome alignments, bottom two blue-(minimap2) and red-(NGMLR) IGV panels). Numerous SNPs in the human *TARDBP* gene (#) are known human polymorphisms present in the targeting construct that differ from the human reference genome, and are only evident as anomalies when aligning to the human reference genome. Two more complex anomalies are denoted by red arrows in intron 2 and intron 5 (region X and Y), which show a number of base positions with high error rates (coloured lines, although the majority of reads map correctly). The intron 5 anomaly additionally shows aberrant sharp changes in coverage. Alignment with an alternative alignment software program, NGMLR (bottom IGV panel, red outline), resolves these latter anomalous features. The only remaining anomalies to account for map to homopolymeric sequences (*h).

**Figure S7. *hFUS* allele theoretical map alignment.** Alignment (minimap2) of *hFUS/hFUS* mouse sequencing reads to a file of the intended humanised allele sequence (green outline top IGV panel). A single imperfect alignment feature (asterisk) maps the correct base in the majority, but is within a repeating element region and shows a higher error rate within reads. Lower four panels show that engineered *loxP* and *FRT* site insertions are only evident as anomalies when aligning to human and mouse reference genomes; bottom and middle right IGV panels are theoretical alignment zooms as indicated; bottom and middle left show the equivalent zoomed regions in human and mouse alignments.

**Figure S8. Regions X and Y in *hTARDBP* introns 2 and 5 precisely map to the boundaries of SINE elements.** Screenshots from UCSC genome browser zoomed in on human introns 2 and 5, with SINE elements indicated by black bars, and regions X and Y indicated by arrows.

**Figure S9. *hTARDBP* allele: C57BL/6 to 129 transition, 5’.** Comparison of C57BL/6J and 129 strain alignments of *hTARDBP/hTARDBP* sequencing reads to reveal the homologous recombination breakpoint in the 5’ homology arm correspondent region, 11 kb upstream of *hTARDBP*. The dotted line marks the equivalent breakpoint position in the two alignments. Some imperfect (but not misaligned) alignment features are also highlighted, with examples given (**). Unannotated regions in the 129 reference genome are also highlighted.

**Figure S10. *hTARDBP* allele: C57BL/6 to 129 transition, 3’.** Comparison of C57BL/6J and 129 strain alignments of *hTARDBP/hTARDBP* sequencing reads to reveal the homologous recombination breakpoint in the 3’ homology arm correspondent region, 5.5 kb downstream of *hTARDBP*. The pair of dotted lines mark the equivalent breakpoint position in the two alignments. Lower panels highlight the breakpoint region at higher resolution. An unannotated region in the 129 reference genome is also highlighted.

**Figure S11. *hFUS* allele: C57BL/6J to 129 transition, 5’.** Comparison of C57BL/6J and 129 strain alignments of *hFUS/hFUS* sequencing reads shows that the proximal ∼30 kb of the 5’ homology arm correspondent region clearly maps precisely to the C57BL/6J reference genome and is derived from the targeting construct. The remaining distal portion of the 5’ homology arm (only the 3’ extremity of this region is shown, marked by C57BL/6J specific insertion, indicated by arrow) has no appreciable divergence between strains to make a clear determination. Some imperfect (but not misaligned) alignment features are also highlighted, with an example given (**). Unannotated regions in the 129 reference genome are also highlighted.

**Figure S12. *hFUS* allele: C57BL/6J to 129 transition, 3’.** Comparison of C57BL/6J and 129 strain alignments of *hFUS/hFUS* sequencing reads shows the 3’ homologous recombination breakpoint can be narrowed to a region 10-15 kb downstream of *hFUS*. The proximal ∼15 kb of the 3’ homology arm correspondent region clearly maps to the C57BL/6J reference genome, with the exception of a single SNP that maps to strain 129 (*129) ∼10 kb downstream of *hFUS*. The remaining distal portion of the 3’ homology arm correspondent region maps to strain 129. Imperfect alignment features are also highlighted.

**Figure S13. *hSOD1* allele: C57BL/6 to 129 transition, 3’.** Comparison of C57BL/6J and 129 strain alignments of *hSOD1/hSOD1* sequencing reads shows the 3’ homologous recombination breakpoint can be narrowed to a region ∼100 kb downstream of *hSOD1*. The proximal ∼100 kb of the 3’ homology arm correspondent region (only the 3’ extremity of this region is shown, marked by C57BL/6 specific insertion, indicated by arrow) clearly maps to the C57BL/6J reference genome, whilst the remaining distal portion of the 3’ homology arm correspondent region maps to strain 129. Unannotated regions in the 129 reference genome are also highlighted.

**Figure S14. Splicing of TDP-43 target exons is unaffected in genomically humanised *hTARDBP* mice.** RT-PCR analysis of *Sort1* exon 18 and *Eif4h* exon 5 inclusion from brain and spinal cord of 14-week old females (n=3 per genotype). No statistically differences (p>0.05) were found for any comparison. Mean ± SD, One-way ANOVA with Dunnett’s post hoc test.

**Figure S15. *<u>hFUS</u>* mRNA and protein expression in 14-week old male tissues.** Expression of *FUS* mRNA and protein in 14-week old male hFUS mice. (A) Quantitative PCR using conserved mouse-human *Fus/FUS* primers was carried out in spinal cord and tibialis anterior (TA) muscle (n=4 per genotype). Data presented as mean ± SD, ns=not statistically significant, *=p≤0.05 calculated using One-way ANOVA with Dunnett’s post hoc test. (B) Immunoblot using pan-mouse-human FUS antibody in male spinal cord and tibialis anterior (TA) muscle tissue (n=4 per genotype per sex). Includes gels used for quantification of FUS protein expression. Data presented as mean ± SD, ns=not statistically significant, calculated using One-way ANOVA with Dunnett’s post hoc test.

**Figure S16. Images of immunoblots used for FUS expression quantification in Figure 6, and for SOD1 expression quantification in figure 4**.

**Figure S17. Modified SHIRPA analyses in *hFUS* mice.** Analysed at 3 and 18 months in *mFus/mFus* and *hFUS/hFUS* mice (3 months: n=17,15,16,13; male *mFus/mFus,* male *hFUS/hFUS,* female *mFus/mFus,* female *hFUS/hFUS,* 18 months: n=12,10,14,11; male *mFus/mFus,* male *hFUS/hFUS,* female *mFus/mFus,* female *hFUS/hFUS*). No significant differences were found for any parameter calculated using Fisher’s exact test or chi square tests.

**Figure S18. Bioinformatics alignment commands.**

**Table S1.**
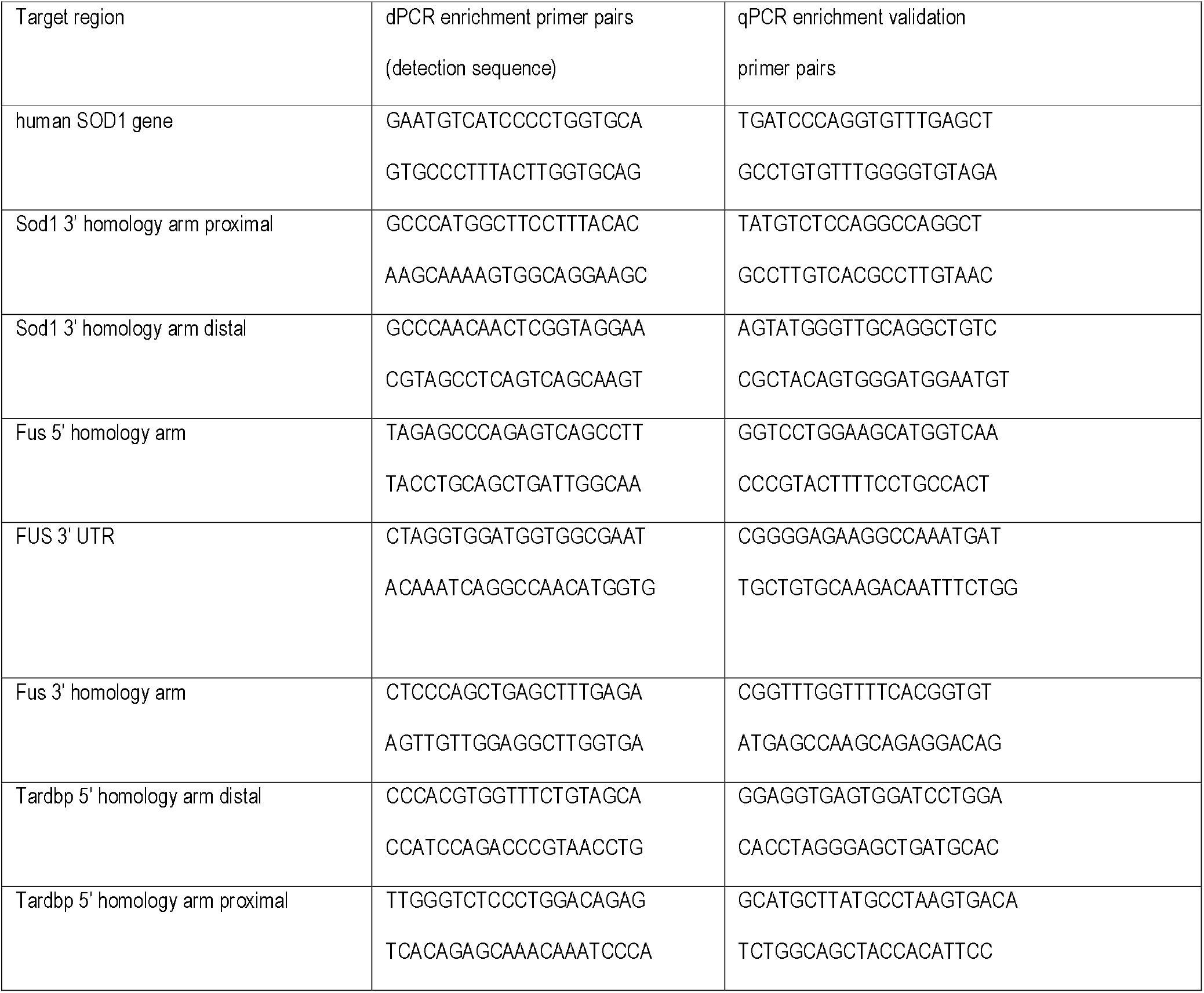
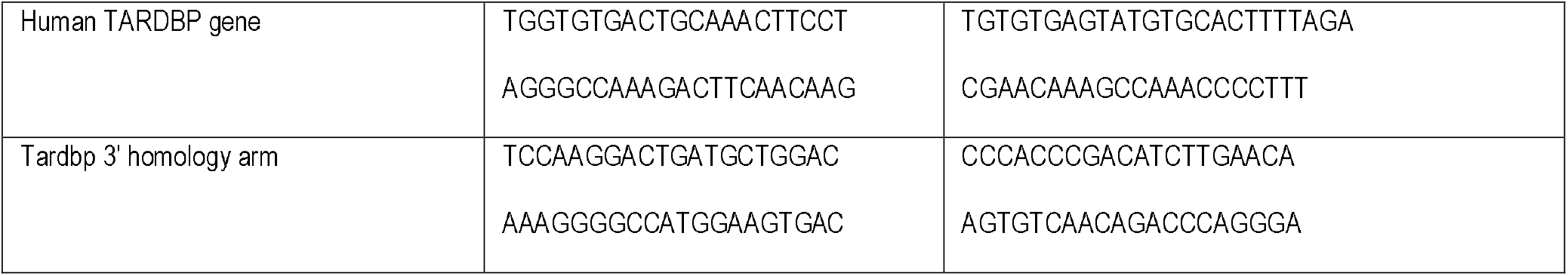
dPCR and qPCR primers used for Xdrop target enrichment and validation.

## Notes

### Competing Interest Statement

The authors have declared no competing interest.

## REFERENCES

Abramzon, Y.A., Fratta, P., Traynor, B.J., and Chia, R. (2020). The Overlapping Genetics of Amyotrophic Lateral Sclerosis and Frontotemporal Dementia. Front Neurosci 14, 42.

Acevedo-Arozena, A., Kalmar, B., Essa, S., Ricketts, T., Joyce, P., Kent, R., Rowe, C., Parker, A., Gray, A., Hafezparast, M., et al. (2011). A comprehensive assessment of the SOD1G93A low-copy transgenic mouse, which models human amyotrophic lateral sclerosis. Dis Model Mech 4, 686–700.

Alexander, G.M., Erwin, K.L., Byers, N., Deitch, J.S., Augelli, B.J., Blankenhorn, E.P., and Heiman-Patterson, T.D. (2004). Effect of transgene copy number on survival in the G93A SOD1 transgenic mouse model of ALS. Brain Res Mol Brain Res 130, 7–15.

Alonso, A., Logroscino, G., Jick, S.S., and Hernan, M.A. (2009). Incidence and lifetime risk of motor neuron disease in the United Kingdom: a population-based study. Eur J Neurol 16, 745–751.

An, H., Rabesahala de Meritens, C., Buchman, V.L., and Shelkovnikova, T.A. (2020). Frameshift peptides alter the properties of truncated FUS proteins in ALS-FUS. Mol Brain 13, 77.

Blondal, T., Gamba, C., Moller Jagd, L., Su, L., Demirov, D., Guo, S., Johnston, C.M., Riising, E.M., Wu, X., Mikkelsen, M.J., et al. (2021). Verification of CRISPR editing and finding transgenic inserts by Xdrop indirect sequence capture followed by short- and long-read sequencing. Methods.

Briese, M., Saal-Bauernschubert, L., Luningschror, P., Moradi, M., Dombert, B., Surrey, V., Appenzeller, S., Deng, C., Jablonka, S., and Sendtner, M. (2020). Loss of Tdp-43 disrupts the axonal transcriptome of motoneurons accompanied by impaired axonal translation and mitochondria function. Acta Neuropathol Commun 8, 116.

Brown, A.-L., Wilkins, O.G., Keuss, M.J., Hill, S.E., Zanovello, M., Lee, W.C., Lee, F.C.Y., Masino, L., Qi, Y.A., Bryce-Smith, S., et al. (2021). Common ALS/FTD risk variants in UNC13A exacerbate its cryptic splicing and loss upon TDP-43 mislocalization. Biorxiv 2021.04.02.438170.

Brown, R.H., and Al-Chalabi, A. (2017). Amyotrophic Lateral Sclerosis. N Engl J Med 377, 162–172.

Codner, G.F., Lindner, L., Caulder, A., Wattenhofer-Donze, M., Radage, A., Mertz, A., Eisenmann, B., Mianne, J., Evans, E.P., Beechey, C.V., et al. (2016). Aneuploidy screening of embryonic stem cell clones by metaphase karyotyping and droplet digital polymerase chain reaction. BMC Cell Biol 17, 30.

Copeland, N.G., Jenkins, N.A., and Court, D.L. (2001). Recombineering: a powerful new tool for mouse functional genomics. Nat Rev Genet 2, 769–779.

Crown, A., McAlary, L., Fagerli, E., Brown, H., Yerbury, J.J., Galaleldeen, A., Cashman, N.R., Borchelt, D.R., and Ayers, J.I. (2020). Tryptophan residue 32 in human Cu-Zn superoxide dismutase modulates prion-like propagation and strain selection. PLoS One 15, e0227655.

de Boer, E.M.J., Orie, V.K., Williams, T., Baker, M.R., De Oliveira, H.M., Polvikoski, T., Silsby, M., Menon, P., van den Bos, M., Halliday, G.M., et al. (2020). TDP-43 proteinopathies: a new wave of neurodegenerative diseases. J Neurol Neurosurg Psychiatry 0, 1–10.

De Giorgio, F., Maduro, C., Fisher, E.M.C., and Acevedo-Arozena, A. (2019). Transgenic and physiological mouse models give insights into different aspects of amyotrophic lateral sclerosis. Dis Model Mech 12, dmm037424.

DeJesus-Hernandez, M., Mackenzie, I.R., Boeve, B.F., Boxer, A.L., Baker, M., Rutherford, N.J., Nicholson, A.M., Finch, N.A., Flynn, H., Adamson, J., et al. (2011). Expanded GGGGCC hexanucleotide repeat in noncoding region of C9ORF72 causes chromosome 9p-linked FTD and ALS. Neuron 72, 245–256.

Devoy, A., Kalmar, B., Stewart, M., Park, H., Burke, B., Noy, S.J., Redhead, Y., Humphrey, J., Lo, K., Jaeger, J., et al. (2017). Humanized mutant FUS drives progressive motor neuron degeneration without aggregation in ‘FUSDelta14’ knockin mice. Brain 140, 2797–2805.

Dib, S., Xiao, S., Miletic, D., and Robertson, J. (2014). Gene targeting of mouse Tardbp negatively affects Masp2 expression. PLoS One 9, e95373.

Dini Modigliani, S., Morlando, M., Errichelli, L., Sabatelli, M., and Bozzoni, I. (2014). An ALS-associated mutation in the FUS 3’-UTR disrupts a microRNA-FUS regulatory circuitry. Nat Commun 5, 4335.

DuVal, M.G., Hinge, V.K., Snyder, N., Kanyo, R., Bratvold, J., Pokrishevsky, E., Cashman, N.R., Blinov, N., Kovalenko, A., and Allison, W.T. (2019). Tryptophan 32 mediates SOD1 toxicity in a in vivo motor neuron model of ALS and is a promising target for small molecule therapeutics. Neurobiol Dis 124, 297–310.

Editorial (2021). Gene therapy needs a long-term approach. Nat Med 27, 563.

Fratta, P., Sivakumar, P., Humphrey, J., Lo, K., Ricketts, T., Oliveira, H., Brito-Armas, J.M., Kalmar, B., Ule, A., Yu, Y., et al. (2018). Mice with endogenous TDP-43 mutations exhibit gain of splicing function and characteristics of amyotrophic lateral sclerosis. EMBO J 37, e98684.

Goodwin, L.O., Splinter, E., Davis, T.L., Urban, R., He, H., Braun, R.E., Chesler, E.J., Kumar, V., van Min, M., Ndukum, J., et al. (2019). Large-scale discovery of mouse transgenic integration sites reveals frequent structural variation and insertional mutagenesis. Genome Res 29, 494–505.

Hardiman, O., Al-Chalabi, A., Chio, A., Corr, E.M., Logroscino, G., Robberecht, W., Shaw, P.J., Simmons, Z., and van den Berg, L.H. (2017). Amyotrophic lateral sclerosis. Nat Rev Dis Primers 3, 17071.

Humphrey, J., Birsa, N., Milioto, C., McLaughlin, M., Ule, A.M., Robaldo, D., Eberle, A.B., Krauchi, R., Bentham, M., Brown, A.L., et al. (2020). FUS ALS-causative mutations impair FUS autoregulation and splicing factor networks through intron retention. Nucleic Acids Res 48, 6889–6905.

Joyce, P.I., McGoldrick, P., Saccon, R.A., Weber, W., Fratta, P., West, S.J., Zhu, N., Carter, S., Phatak, V., Stewart, M., et al. (2015). A novel SOD1-ALS mutation separates central and peripheral effects of mutant SOD1 toxicity. Hum Mol Genet 24, 1883–1897.

Klim, J.R., Williams, L.A., Limone, F., Guerra San Juan, I., Davis-Dusenbery, B.N., Mordes, D.A., Burberry, A., Steinbaugh, M.J., Gamage, K.K., Kirchner, R., et al. (2019). ALS-implicated protein TDP-43 sustains levels of STMN2, a mediator of motor neuron growth and repair. Nat Neurosci 22, 167–179.

Kwiatkowski, T.J., Jr., Bosco, D.A., Leclerc, A.L., Tamrazian, E., Vanderburg, C.R., Russ, C., Davis, A., Gilchrist, J., Kasarskis, E.J., Munsat, T., et al. (2009). Mutations in the FUS/TLS gene on chromosome 16 cause familial amyotrophic lateral sclerosis. Science 323, 1205–1208.

Kwon, D. (2021). Failure of genetic therapies for Huntington’s devastates community. Nature.

Lee, Y., and Rio, D.C. (2015). Mechanisms and Regulation of Alternative Pre-mRNA Splicing. Annu Rev Biochem 84, 291–323.

Li, H. (2018). Minimap2: pairwise alignment for nucleotide sequences. Bioinformatics 34, 3094–3100.

Ling, S.C., Dastidar, S.G., Tokunaga, S., Ho, W.Y., Lim, K., Ilieva, H., Parone, P.A., Tyan, S.H., Tse, T.M., Chang, J.C., et al. (2019). Overriding FUS autoregulation in mice triggers gain-of-toxic dysfunctions in RNA metabolism and autophagy-lysosome axis. Elife 8, e40811.

Lopez-Erauskin, J., Tadokoro, T., Baughn, M.W., Myers, B., McAlonis-Downes, M., Chillon-Marinas, C., Asiaban, J.N., Artates, J., Bui, A.T., Vetto, A.P., et al. (2018). ALS/FTD-Linked Mutation in FUS Suppresses Intra-axonal Protein Synthesis and Drives Disease Without Nuclear Loss-of-Function of FUS. Neuron 100, 816–830 e817.

Ma, X.R., Prudencio, M., Koike, Y., Vatsavayai, S.C., Kim, G., Harbinski, F., Rodriguez, C.M., Schmidt, H.B., Cummings, B.B., Wyatt, D.W., et al. (2021). TDP-43 represses cryptic exon inclusion in FTD/ALS gene UNC13A. Biorxiv 2021.*04.02.438213*.

Mackenzie, I.R., Bigio, E.H., Ince, P.G., Geser, F., Neumann, M., Cairns, N.J., Kwong, L.K., Forman, M.S., Ravits, J., Stewart, H., et al. (2007). Pathological TDP-43 distinguishes sporadic amyotrophic lateral sclerosis from amyotrophic lateral sclerosis with SOD1 mutations. Ann Neurol 61, 427–434.

Mandillo, S., Heise, I., Garbugino, L., Tocchini-Valentini, G.P., Giuliani, A., Wells, S., and Nolan, P.M. (2014). Early motor deficits in mouse disease models are reliably uncovered using an automated home-cage wheel-running system: a cross-laboratory validation. Dis Model Mech 7, 397–407.

Mejzini, R., Flynn, L.L., Pitout, I.L., Fletcher, S., Wilton, S.D., and Akkari, P.A. (2019). ALS Genetics, Mechanisms, and Therapeutics: Where Are We Now? Front Neurosci 13, 1310.

Melamed, Z., Lopez-Erauskin, J., Baughn, M.W., Zhang, O., Drenner, K., Sun, Y., Freyermuth, F., McMahon, M.A., Beccari, M.S., Artates, J.W., et al. (2019). Premature polyadenylation-mediated loss of stathmin-2 is a hallmark of TDP-43-dependent neurodegeneration. Nat Neurosci 22, 180–190.

Mitchell, J.C., McGoldrick, P., Vance, C., Hortobagyi, T., Sreedharan, J., Rogelj, B., Tudor, E.L., Smith, B.N., Klasen, C., Miller, C.C., et al. (2013). Overexpression of human wild-type FUS causes progressive motor neuron degeneration in an age- and dose-dependent fashion. Acta Neuropathol 125, 273–288.

Nagano, S., Takahashi, Y., Yamamoto, K., Masutani, H., Fujiwara, N., Urushitani, M., and Araki, T. (2015). A cysteine residue affects the conformational state and neuronal toxicity of mutant SOD1 in mice: relevance to the pathogenesis of ALS. Hum Mol Genet 24, 3427–3439.

Nair, R.R., Corrochano, S., Gasco, S., Tibbit, C., Thompson, D., Maduro, C., Ali, Z., Fratta, P., Arozena, A.A., Cunningham, T.J., et al. (2019). Uses for humanised mouse models in precision medicine for neurodegenerative disease. Mamm Genome 30, 173–191.

Nelson, P.T., Dickson, D.W., Trojanowski, J.Q., Jack, C.R., Boyle, P.A., Arfanakis, K., Rademakers, R., Alafuzoff, I., Attems, J., Brayne, C., et al. (2019). Limbic-predominant age-related TDP-43 encephalopathy (LATE): consensus working group report. Brain 142, 1503–1527.

Neumann, M., Rademakers, R., Roeber, S., Baker, M., Kretzschmar, H.A., and Mackenzie, I.R. (2009). A new subtype of frontotemporal lobar degeneration with FUS pathology. Brain 132, 2922–2931.

Perri, E.R., Parakh, S., Vidal, M., Mehta, P., Ma, Y., Walker, A.K., and Atkin, J.D. (2020). The Cysteine (Cys) Residues Cys-6 and Cys-111 in Mutant Superoxide Dismutase 1 (SOD1) A4V Are Required for Induction of Endoplasmic Reticulum Stress in Amyotrophic Lateral Sclerosis. J Mol Neurosci 70, 1357–1368.

Picher-Martel, V., Brunet, F., Dupre, N., and Chrestian, N. (2020). The Occurrence of FUS Mutations in Pediatric Amyotrophic Lateral Sclerosis: A Case Report and Review of the Literature. J Child Neurol 35, 556–562.

Renton, A.E., Majounie, E., Waite, A., Simon-Sanchez, J., Rollinson, S., Gibbs, J.R., Schymick, J.C., Laaksovirta, H., van Swieten, J.C., Myllykangas, L., et al. (2011). A hexanucleotide repeat expansion in C9ORF72 is the cause of chromosome 9p21-linked ALS-FTD. Neuron 72, 257–268.

Rogers, D.C., Fisher, E.M., Brown, S.D., Peters, J., Hunter, A.J., and Martin, J.E. (1997). Behavioral and functional analysis of mouse phenotype: SHIRPA, a proposed protocol for comprehensive phenotype assessment. Mamm Genome 8, 711–713.

Rosen, D.R., Siddique, T., Patterson, D., Figlewicz, D.A., Sapp, P., Hentati, A., Donaldson, D., Goto, J., O’Regan, J.P., Deng, H.X., et al. (1993). Mutations in Cu/Zn superoxide dismutase gene are associated with familial amyotrophic lateral sclerosis. Nature 362, 59–62.

Sabatelli, M., Moncada, A., Conte, A., Lattante, S., Marangi, G., Luigetti, M., Lucchini, M., Mirabella, M., Romano, A., Del Grande, A., et al. (2013). Mutations in the 3’ untranslated region of FUS causing FUS overexpression are associated with amyotrophic lateral sclerosis. Hum Mol Genet 22, 4748–4755.

Saccon, R.A., Bunton-Stasyshyn, R.K., Fisher, E.M., and Fratta, P. (2013). Is SOD1 loss of function involved in amyotrophic lateral sclerosis? Brain 136, 2342–2358.

Sedlazeck, F.J., Rescheneder, P., Smolka, M., Fang, H., Nattestad, M., von Haeseler, A., and Schatz, M.C. (2018). Accurate detection of complex structural variations using single-molecule sequencing. Nat Methods 15, 461–468.

Stewart, M., Lau, P., Banks, G., Bains, R.S., Castroflorio, E., Oliver, P.L., Dixon, C.L., Kruer, M.C., Kullmann, D.M., Acevedo-Arozena, A., et al. (2019). Loss of Frrs1l disrupts synaptic AMPA receptor function, and results in neurodevelopmental, motor, cognitive and electrographical abnormalities. Dis Model Mech 12.

Suk, T.R., and Rousseaux, M.W.C. (2020). The role of TDP-43 mislocalization in amyotrophic lateral sclerosis. Mol Neurodegener 15, 45.

Taylor, J.P., Brown, R.H., Jr., and Cleveland, D.W. (2016). Decoding ALS: from genes to mechanism. Nature 539, 197–206.

Urwin, H., Josephs, K.A., Rohrer, J.D., Mackenzie, I.R., Neumann, M., Authier, A., Seelaar, H., Van Swieten, J.C., Brown, J.M., Johannsen, P., et al. (2010). FUS pathology defines the majority of tau- and TDP-43-negative frontotemporal lobar degeneration. Acta Neuropathol 120, 33–41.

Valenzuela, D.M., Murphy, A.J., Frendewey, D., Gale, N.W., Economides, A.N., Auerbach, W., Poueymirou, W.T., Adams, N.C., Rojas, J., Yasenchak, J., et al. (2003). High-throughput engineering of the mouse genome coupled with high-resolution expression analysis. Nature biotechnology 21, 652–659.

Vance, C., Rogelj, B., Hortobagyi, T., De Vos, K.J., Nishimura, A.L., Sreedharan, J., Hu, X., Smith, B., Ruddy, D., Wright, P., et al. (2009). Mutations in FUS, an RNA processing protein, cause familial amyotrophic lateral sclerosis type 6. Science 323, 1208–1211.

Wils, H., Kleinberger, G., Janssens, J., Pereson, S., Joris, G., Cuijt, I., Smits, V., Ceuterick-de Groote, C., Van Broeckhoven, C., and Kumar-Singh, S. (2010). TDP-43 transgenic mice develop spastic paralysis and neuronal inclusions characteristic of ALS and frontotemporal lobar degeneration. Proc Natl Acad Sci U S A 107, 3858–3863.

Xu, Y.F., Gendron, T.F., Zhang, Y.J., Lin, W.L., D’Alton, S., Sheng, H., Casey, M.C., Tong, J., Knight, J., Yu, X., et al. (2010). Wild-type human TDP-43 expression causes TDP-43 phosphorylation, mitochondrial aggregation, motor deficits, and early mortality in transgenic mice. J Neurosci 30, 10851–10859.

Zhao, M., Kim, J.R., van Bruggen, R., and Park, J. (2018). RNA-Binding Proteins in Amyotrophic Lateral Sclerosis. Mol Cells 41, 818–829.

Zhu, F., Nair, R.R., Fisher, E.M.C., and Cunningham, T.J. (2019). Humanising the mouse genome piece by piece. Nat Commun 10, 1845.

